# Multiple Mechanisms Underlie State-Independent Inhibitory Effects of Norfluoxetine on TREK-2 K2P Channels

**DOI:** 10.1101/2020.10.29.360966

**Authors:** Peter Proks, Marcus Schewe, Linus J. Conrad, Shanlin Rao, Kristin Rathje, Karin E. J. Rödström, Elisabeth P. Carpenter, Thomas Baukrowitz, Stephen J Tucker

## Abstract

The TREK subfamily of Two-Pore Domain (K2P) K^+^ channels are inhibited by fluoxetine and its metabolite, norfluoxetine (NFx). Although not the principal targets of this antidepressant, TREK channel inhibition by NFx has provided important insights into the conformational changes associated with channel gating and highlighted the role of the selectivity filter in this process. But despite the availability of TREK-2 crystal structures with NFx bound, the precise mechanisms underlying NFx inhibition remain elusive. NFx has previously been proposed to be a state-dependent inhibitor, but its binding site suggests many possible ways in which this positively charged drug might inhibit channel activity. Here we show that NFx exerts multiple effects on single channel behavior that influence both the open and closed states of the channel, and that the channel can become highly activated by 2-APB whilst remaining in the down conformation. We also show that that the inhibitory effects of NFx are unrelated to its positive charge, but can be influenced by agonists such as ML335 which alter filter stability, as well as by an intrinsic voltage-dependent gating process within the filter. NFx therefore not only inhibits channel activity by altering the equilibrium between up and down conformations, but can also directly influence filter gating. These results provide further insight into the complex allosteric mechanisms that modulate filter-gating in TREK K2P channels and highlight the different ways that filter gating can be regulated to permit polymodal regulation.

## Introduction

Within the Two-Pore domain (K2P) family of K^+^ channels, the TREK subfamily (K2P2.1/TREK-1, K2P10.1/TREK-2, and K2P4.1/TRAAK) exhibit polymodal regulation by diverse chemical and physical stimuli that couple many different cellular and environmental signals to changes in cellular electrical activity (Enyedi and Czirjak, 2010; Niemeyer et al., 2016). TREK channels are located throughout the central and peripheral nervous system where they are involved in variety of processes including mechanosensation, thermosensation and nociception (Djillani et al., 2019a). As a consequence of their apparent role in these tissues, selective TREK channel agonists have been proposed as potential analgesics (Mathie and Veale, 2015; Vivier et al., 2016) and several inhibitors are also considered possible antidepressants (Heurteaux et al., 2006; Djillani et al., 2019b). Understanding the mechanisms by which such small molecules and other compounds modulate TREK channel activity is therefore important to fully realize their therapeutic potential.

In a previous study we solved crystal structures of the human TREK-2 channel in two distinct structural states known as the ‘up’ and ‘down’ conformations (Dong et al., 2015). In that same study, we also determined structures of TREK-2 in complex with two known inhibitors, namely fluoxetine and its active metabolite, norfluoxetine (NFx). Fluoxetine (Prozac™) is a commonly prescribed antidepressant and although its principal action as a selective serotonin reuptake inhibitor is well characterized, its inhibitory effects on TREK channels remain of interest due to the reported link between TREK-1 and depression and the fact it is one of few relatively high-affinity blockers of TREK channels currently available (Kennard et al., 2005; Heurteaux et al., 2006). Furthermore, the crystal structures of TREK-2 revealed that NFx binds in the inner cavity of TREK-2 within side fenestrations formed by a gap between the transmembrane (TM) domains. Consequently, the NFx binding site is only available in the down state because these gaps are not present in the up state. However, the relative activity of these up and down conformations is unknown and the precise mechanism by which NFx binding leads to inhibition of TREK channel activity remains unclear.

It has recently been shown that some K2P channels possess a lower gate analogous to the classical helix bundle-crossing found in many other types of K^+^ channels (Li et al., 2020; Rödstrom et al., 2020). However, most K2P channels, including TREK channels, do not have a lower gate (Brohawn et al., 2012; Miller and Long, 2012; Lolicato et al., 2014; Dong et al., 2015). Instead they appear to gate primarily within their selectivity filter (Zilberberg et al., 2001; Bagriantsev et al., 2011; Piechotta et al., 2011; Schewe et al., 2016; Nematian-Ardestani et al., 2020) and current models for TREK channel gating propose that movement of the TM helices can regulate this filter gating mechanism but do not constrict enough at this lower region to prevent K^+^ permeation (Brohawn et al., 2012; Lolicato et al., 2014; Dong et al., 2015).

This current model for TREK channel gating is shown in **Figure 1A**. The two global states of the channel, the up and down conformations, are thought to regulate TREK channel activity by controlling the dynamics of the filter gate. Initially, when these two conformations were identified it was thought that NFx binding to the down state would not only stabilize that conformation, but also impair movement to the up conformation thereby representing a possible mechanism for the apparent state-dependent effects of NFx on TREK channel activity (Kennard et al., 2005). However, it was later shown that when the filter gate was activated directly, either by mutations near the filter, or by using Rb^+^ as the permeant ion, that NFx inhibition was not affected (McClenaghan et al., 2016). Furthermore, it was shown that BL1249 directly activates the filter gate when the channel is in the down state (Schewe et al., 2019). These observations imply the filter gate can be conductive when the channel is in the down conformation and that movement to the up state is not required for the filter gate to open. The model also proposed that movement of the TM-helices modulates the relative activity of the filter gate with the up conformation enabling a higher open probability when the up state is stabilized by physiological stimuli e.g. membrane stretch (Aryal et al., 2017; Clausen et al., 2017).

**Figure 1:**
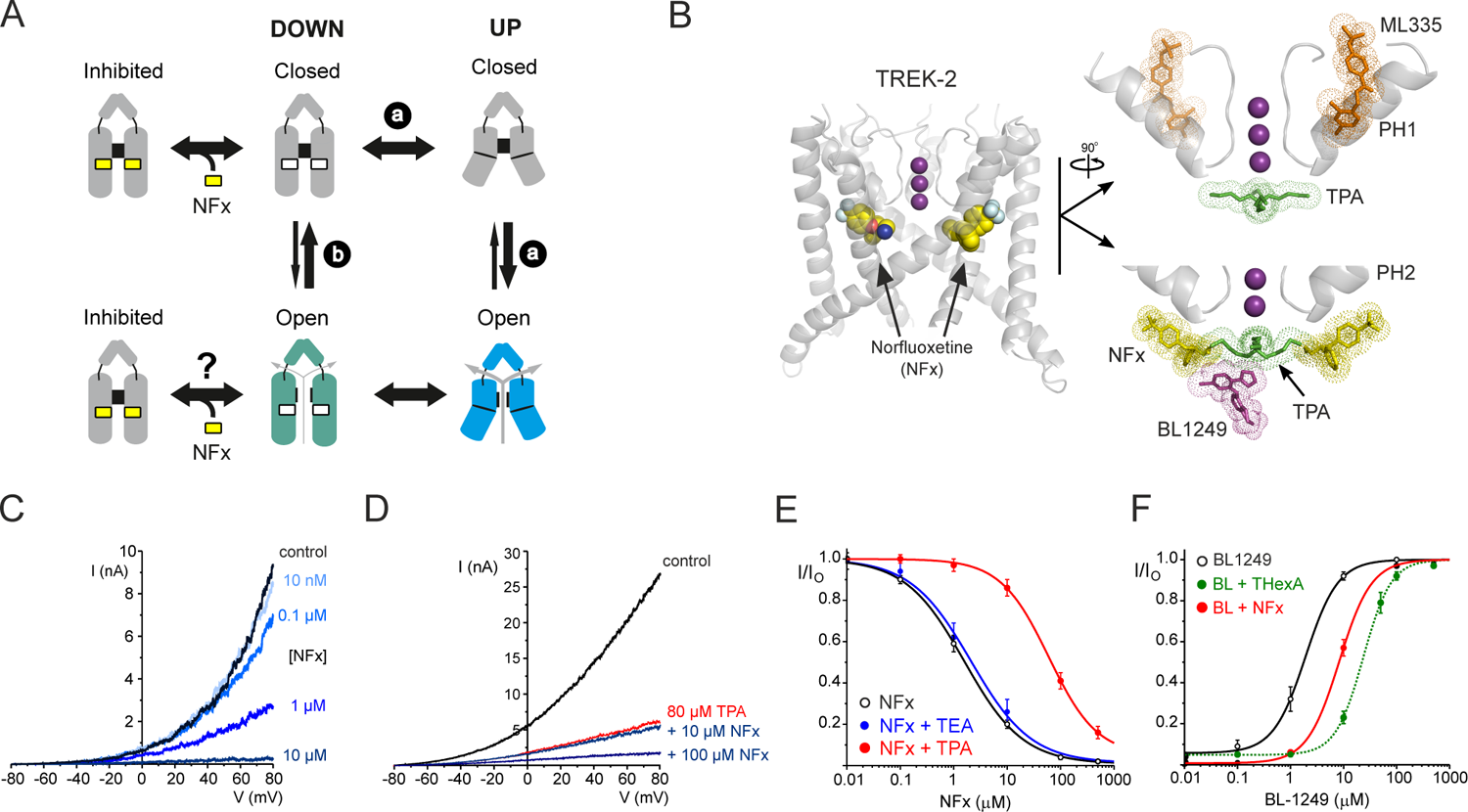
The current model for TREK channel gating and NFx binding sites. A. The TM helices exist in two states (up and down) but it is unclear whether opening of the filter gate requires movement to the up conformation (via route a), whether it can open independently in the down state (via route b), or even whether both options are possible. Current models also suggest that openings from the down state may result in a lower activity channel than when it is in the up state because many activatory mechanisms, e.g. membrane stretch promote movement to the up state. Binding sites for norfluoxetine (NFx) do not exist in the up state and NFx binding will alter the equilibrium between these two conformations of the TM-helices but is unclear whether NFx binding is state-dependent and only stabilizes the closed state of the channel. The presence of positively charged NFx bound within the inner pore may also cause direct pore block and/or allosteric effects on the filter gating mechanism itself. **B**. Left: a view of the structure of TREK-2 in the down state showing NFx (as vdW spheres) bound within the fenestrations (PDB ID: 4XDK). K^+^ ions in the filter are shown as purple spheres. Right: expanded views of other drugs binding sites near the filter. The top panel (rotated by 90°) shows Pore-Helix 1 (PH1) and the position of the ML335 (orange) which does not overlap with that of TPA (green) below the filter). In the bottom panel is the inner cavity below the filter showing the position of NFx (yellow), TPA (green) and BL1249 (purple) when bound to channel. The position of Pore-Helix 2 (PH2) is also shown. The binding sites for all three ligands are in close proximity and exhibit partial overlap but not with ML335. **C.** Representative traces of macroscopic TREK-2 currents elicited by voltage ramps between −80 and +80mV in giant excised patches from *Xenopus* oocytes measured in control solution and various bath concentrations of NFx, as indicated. **D**. Similar representative traces of macroscopic TREK-2 currents showing reduced inhibition by NFx in the presence of 80 µM TPA. Block by 80 µM TPA alone shown in red. **E.** NFx inhibition of TREK-2 currents at +40 mV in *Xenopus* oocytes on its own (*IC_50_*=2.7 μM; *h*=1.0 n=19) and in the presence of 100 mM TEA (*IC_50_*=3.8 μM; *h*=0.8 n=7) or 80μM TPA (*IC_50_*=65μM; *h*=1.2 *a*=0.05; n=12), as indicated. **F.** BL1249 activation of TREK-2 currents in *Xenopus* oocytes on its own (*IC_50_*=2.5μM; *h*=1.9, n=13) or in the presence of NFx. For comparison the previously reported shift in the presence of 5μM THexA (Schewe et al., 2019) is also shown as a dotted green line.

However, this model is based on ensemble measurements of channel activity where it remains difficult to exclude the possibility that openings of the filter gate from a ‘lower activity’ down state remain coupled to brief structural movements of the TM-helices into the up conformation. The true state-dependence of NFx inhibition therefore remains unknown. Also such macroscopic measurements cannot determine conclusively whether the channel can become fully activated whilst remaining in the down conformation.

Other mechanisms of inhibition might also contribute to its effects on channel activity. For example, in addition to any effect on the equilibrium between up and down conformations, the binding of NFx within the inner pore (**Figure 1B**) and its intrinsic positive charge suggests that it could also directly block or impair K^+^ permeation through the inner cavity. Finally, the proximity of the NFx binding sites to the filter also suggests it might exert additional allosteric effects on the filter gate itself.

The role of these different possible mechanisms of inhibition by NFx have not been fully explored and their relative contribution remains unknown. Understanding these processes is not only important for dissecting the mechanism of NFx inhibition, but also in determining how filter gating is coupled to the different conformational states of the TM-helices. In this study we have therefore examined the inhibition of TREK-2 channel activity by NFx at both the macroscopic and single channel level. Our results provide new insight into the state-independent inhibitory effects of NFx on both the open and closed states of the channel, and we show that the channel can become highly activate when it is the down conformation.

## Results and Discussion

### Interactions between agonists and inhibitors within the inner cavity

To begin to dissect its inhibitory effects, we first examined the interaction between NFx and other ligands which also modulate channel activity. Quaternary ammonium (QA) ions such as Tetrapentylammonium (TPA) are known to bind to a variety of K^+^ channels deep within the cavity just below the selectivity filter, and have proven useful tools of in the study of channel pore structure and gating (Armstrong, 1971; Baukrowitz and Yellen, 1996; Piechotta et al., 2011). In TREK-2, the two binding sites for NFx are also located below the selectivity filter and though not directly below the entrance to the filter, these sites are close enough to partially overlap with the central binding site for TPA (**Figure 1B**) (Rapedius et al., 2012; Dong et al., 2015) and so might be expected to result in interactions between these inhibitors; we therefore examined whether NFx inhibition of TREK-2 was affected by the presence of TPA.

As reported previously, NFx itself produces concentration-dependent inhibitory effects in giant inside-out (**Figure 1C**). The *IC_50_* for channel inhibition was ∼3 μM at physiological pH7.4. However, when similar dose-response curves were measured in the presence of 80 μM TPA there was a marked reduction in the efficacy of NFx inhibition (*IC_50_* to ∼65 µM; **Figure 1D,E**). Importantly, there was little shift in NFx inhibition with the smaller sized QA, Tetraelthylammonium (TEA) which is not predicted to overlap with the NFx binding site (**Figure 1E**). A recent crystal structure also revealed a binding site for BL1249, a TREK channel activator, which partially overlaps with the QA binding site (**Figures 1B**) and that BL1249 activation is affected by the presence of Tetrahexylammonium (THexA) (Schewe et al., 2019). Consistent with the close proximity of these three ligand binding sites, we also observed a reduced activatory effect of BL1249 in the presence NFx (**Figure 1F**).

Recent crystal structures show that another TREK-2 agonist, ML335 binds to an unrelated site behind the selectivity filter that does not overlap with the NFx binding site (Lolicato et al., 2017) (**Figure 1B**). Interestingly, we still observed a marked reduction in NFx inhibition in the presence of 50 µM ML335 (*IC_50_* increased from ∼3 to ∼160 μM; **Figure 2A,C**). This effect appears specific because activation by another agonist, 2-aminoethoxydiphenyl borate (2-APB), whose binding site is predicted to be in the C-terminus (Zhuo et al., 2015) had little effect on NFx inhibition (**Figure 2B,C**). Importantly, the channel retains this sensitivity to NFx even when activated by 2-APB thereby confirming that the NFx-sensitive down conformation can adopt a high activity gating mode.

**Figure 2:**
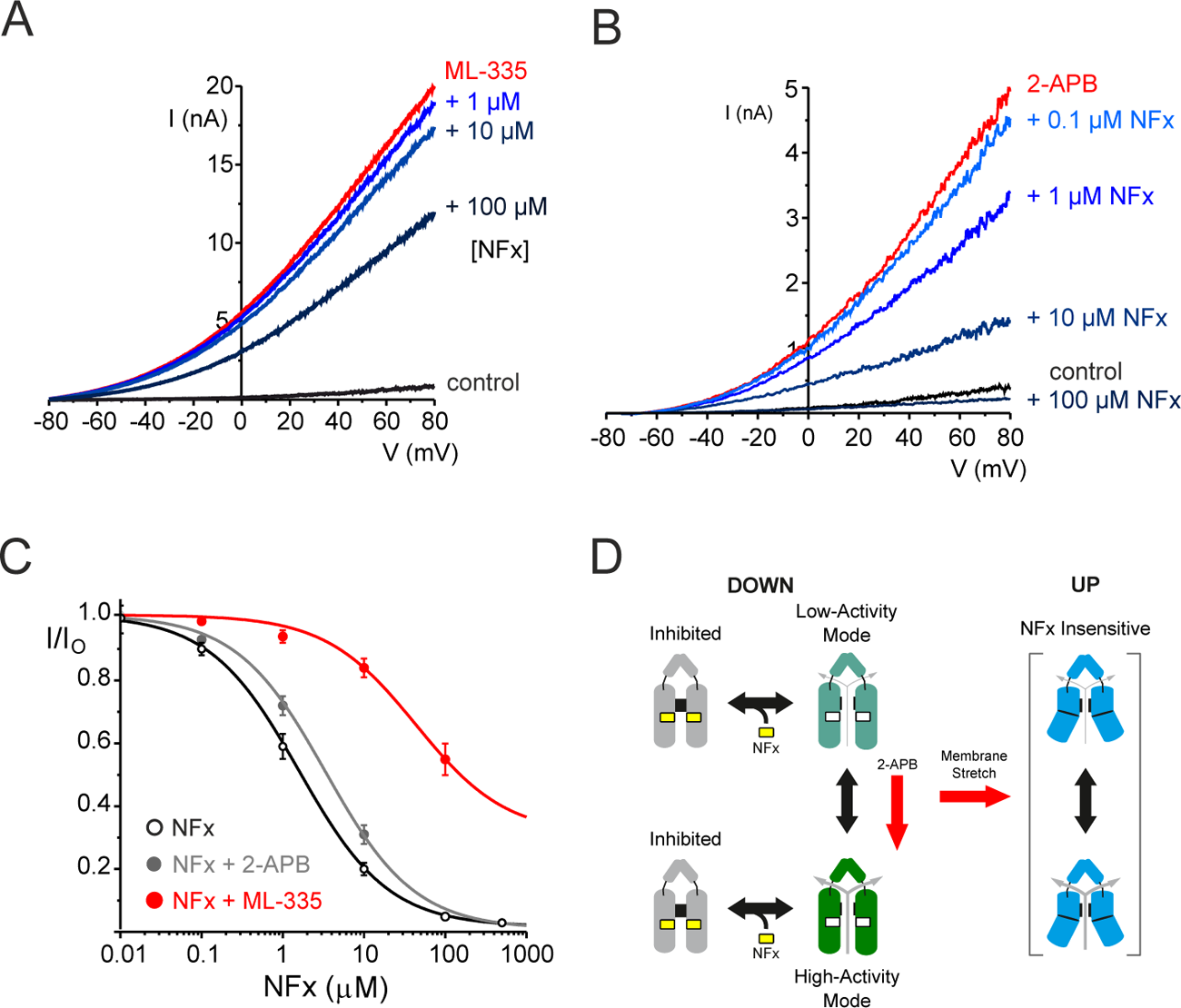
Direct and allosteric interactions of NFx with TREK-2. **A**. Representative traces of macroscopic TREK-2 currents elicited by voltage ramps between −80 and +80mV in giant excised patches from *Xenopus* oocytes measured in control solution, in the presence of 50μM ML-335 alone and with various added bath concentrations of NFx, as indicated. **B**. Similar representative traces of TREK-2 currents in the absence (control) or presence of the activator, 1 mM 2-APB alone, and with added concentrations of NFx, as indicated. **C**. Dose response curves for NFx inhibition of TREK-2 currents on their own (*IC_50_*=2.7μM; *h*=1.7 n=19) or in the presence of 1mM 2-APB (*IC_50_*=3.8μM; *h*=0.6 n=11) or 50μM ML-335 (*IC_50_*=164μM; *h*=0.8; n=7), as indicated. Note the large shift in NFx sensitivity that results from ML335 activation, but not 2-APB activation. **D.** Modified gating cartoon indicating gating modes with different activities rather than distinct open/closed states. The red arrow shows that 2-APB promotes a highly active state with unaltered NFx sensitivity suggesting NFx inhibition is not state-dependent. Other factors such as membrane stretch promote formation of various NFx-insensitive up conformations.

It has recently been demonstrated that ML335 binding behind the selectivity filter directly activates the filter gating mechanism by affecting the dynamic flexibility of the loops and pore helices supporting the selectivity filter (Lolicato et al., 2020). The antagonistic effect of ML335 we observe on NFx inhibition from a non-overlapping site strongly suggests that NFx may inhibit TREK-2 via allosteric effects on the filter gating mechanism itself. Furthermore, our results indicate that the mechanism of activation by 2-APB must be different because NFx sensitivity is not altered by 2-APB and so the channel must remain in the down conformation when activated by 2-APB.

The original gating cartoon in **Figure 1B** is therefore better represented by ‘modes’ of gating capable of supporting different activities rather than distinct open/closed states and this is now shown in **Figure 2D**. However, from these macroscopic recordings alone, NFx provides little information about possible up conformations as they are NFx-insensitive and it is difficult to exclude other more direct effects of NFx on K^+^ permeation, or whether it preferentially binds to the closed state of the channel. We therefore examined the effects of NFx inhibition on the behavior of TREK-2 at the single channel level.

### Characterization of TREK-2 single channel behavior

Detailed analysis of the effect of drugs on single channel behavior can provide important insights into the mechanism of drug action. In particular if NFx acts as a state-dependent blocker that only affects the closed state then such an effect should be evident from these recordings. However, there are two major problems when attempting to study the behavior of wild-type TREK2 and its inhibition by NFx. The first problem comes from the variable kinetics and conductances reported for wild-type TREK-2 single channels (Kang et al., 2007). This is thought to arise from the multiple isoforms produced by alternative translation initiation sites within the N-terminus (Simkin et al., 2008), but irrespective of the cause, these variations complicates the analysis of single channel data. The second issue is that, unless activated, individual TREK-2 channels have a very low ‘resting’ open probability (*P_o_*) which makes detailed analysis of the effects of an inhibitor extremely challenging.

In a previous study we measured the activity of purified TREK-2 channels reconstituted into a lipid bilayer (Clausen et al., 2017). These purified proteins were the same as those used to obtain crystal structures of TREK-2 with NFx bound (Dong et al., 2015), and although truncated at both the N- and C-termini they still produce functional channels that can be inhibited by NFx and activated by BL1249 (Dong et al., 2015). Similar truncations in TREK-1 also retain their activation by ML335 (Lolicato et al., 2017). Furthermore, when measured in bilayers these truncated proteins do not produce the highly variable single channel conductances that WT TREK2 exhibits when expressed in heterologous systems. We therefore chose to examine the effects of NFx on recordings of single TREK-2 channels in this bilayer system.

### Characterization of single TREK-2 channels in lipid bilayers

Regardless of their orientation in the bilayer, we found the *P_o_* of most reconstituted TREK-2 channels was strongly voltage-dependent with inward currents having a much lower *P_o_* than outward currents (**Supplementary Figure 1A,B**). This “standard” behavior resulted in outwardly-rectifying macroscopic currents similar to that observed in many previous recordings of WT TREK-2 currents expressed in heterologous systems, but the *P_o_* of these channels was not stationary over long periods of time meaning that a detailed analysis of their inhibition by NFx would be difficult.

However, in ∼10% of recordings, we observed a high *P_O_* mode of behavior for both outward and inward currents which resulted in a quasi-symmetrical current-voltage relationship (**Supplementary Figure 1C,D**). Interestingly, if several channels were present in a recording, they would all exhibit either the standard or high *P_o_* mode of gating but these different modes were never observed together. The reasons underlying this high *P_o_* mode and their conformational identity are uncertain yet they are unlikely to be predominantly in the up conformation as they remain NFx sensitive. However, the stability of their single channel behavior over long periods makes them particularly suitable for analyzing the inhibitory effects of NFx.

We therefore examined the kinetics of these channels in the absence of NFx; the distributions of openings of both inward (−60 mV) and outward (+60 mV) currents in both standard and high *P_o_* mode were well-fitted by a single exponential (**Supplementary Figure S2** and **Supplementary Table S1**). The distribution of closings of both inward and outward TREK-2 channels in the standard mode were well fit by five exponentials (**Supplementary Figure S2**), but only the shortest two of these exponential components were present in the high *P_o_* mode (**Supplementary Figure S2B**).

**Table 1.** Comparison of kinetic parameters of single TREK-2 channel recording depicted and analyzed in. Figures 4 and Supplementary Figure S1 **& S5.** τ_O_, mean open time; τ_Ci_ and A_Ci_, mean lifetimes and corresponding areas of closed states (*i*=1-5).

| [NFx](μM) | V (mV) | $\tau_o$ (ms) | $\tau_{C1}$ (ms) | $A_{C1}$ (%) | $\tau_{C2}$ (ms) | $A_{C2}$ (%) | $\tau_{C3}$ (ms) | $A_{C3}$ (%) | $\tau_{C4}$ (ms) | $A_{C4}$ (%) | $\tau_{C5}$ (ms) | $A_{C5}$ (%) |
| --- | --- | --- | --- | --- | --- | --- | --- | --- | --- | --- | --- | --- |
| 0 | -100 | 0.17 | 0.107 | 97 | 0.56 | 3 | - | - | - | - | - | - |
| 0 | -80 | 0.25 | 0.086 | 97 | 0.68 | 3 | - | - | - | - | - | - |
| 0 | -60 | 0.37 | 0.073 | 96 | 0.81 | 4 | - | - | - | - | - | - |
| 0 | -40 | 0.58 | 0.058 | 96 | 0.88 | 4 | - | - | - | - | - | - |
| 0 | -20 | 0.88 | 0.049 | 92 | 0.95 | 8 | - | - | - | - | - | - |
| 0 | 20 | 3.2 | 0.086 | 74 | 1.01 | 26 | - | - | - | - | - | - |
| 0 | 40 | 4.7 | 0.097 | 72 | 1.06 | 28 | - | - | - | - | - | - |
| 0 | 60 | 4.6 | 0.102 | 71 | 1.09 | 29 | - | - | - | - | - | - |
| 0 | 80 | 3.8 | 0.107 | 74 | 1.07 | 26 | - | - | - | - | - | - |
| 0 | 100 | 2.7 | 0.106 | 77 | 1.10 | 23 | - | - | - | - | - | - |
| 100 | -100 | 0.20 | 0.110 | 88 | 0.37 | 8.0 | 3.0 | 0.6 | 40 | 2.2 | 140 | 1.0 |
| 100 | -80 | 0.27 | 0.110 | 89 | 0.37 | 6.0 | 9.5 | 1.0 | 58 | 2.8 | 140 | 0.8 |
| 100 | -60 | 0.35 | 0.082 | 85 | 0.30 | 9.5 | 1.6 | 1.2 | 44 | 3.4 | 230 | 0.6 |
| 100 | -40 | 0.51 | 0.069 | 85 | 0.35 | 9.3 | 2.5 | 1.7 | 52 | 4.2 | 520 | 0.2 |
| 100 | -20 | 0.54 | 0.061 | 81 | 0.35 | 10.7 | 2.5 | 2.5 | 45 | 5.0 | 260 | 0.6 |
| 100 | 20 | 0.53 | 0.041 | 74 | 0.30 | 15.5 | 1.3 | 4.4 | 23 | 4.9 | 90 | 1.5 |
| 100 | 40 | 0.69 | 0.083 | 75 | 0.36 | 13.7 | 4.8 | 3.3 | 39 | 6.2 | 160 | 2.0 |
| 100 | 60 | 0.60 | 0.059 | 75 | 0.31 | 13.9 | 2.1 | 3.5 | 41 | 5.7 | 160 | 1.6 |
| 100 | 80 | 0.49 | 0.055 | 76 | 0.32 | 11.9 | 2.9 | 1.9 | 78 | 7.7 | 490 | 2.6 |
| 100 | 100 | 0.44 | 0.064 | 77 | 0.30 | 9.2 | 1.9 | 2.5 | 72 | 7.5 | 220 | 4.1 |

### The effect of NFx on the properties of single TREK-2 channels

We next examined the inhibitory effects of NFx on the well-behaved kinetics of TREK-2 in this high *P_o_* mode. As a control for any indirect effects of NFx on the properties of the bilayer (Kapoor et al., 2019), we first tested whether high concentrations of NFx could affect the elastic modulus of the DPhPC bilayers used in our experiments using the electrostriction method (Vitovic et al., 2013). However, we found no obvious effect of 1-1000 μM NFx on the modulus of elasticity (*E*_Ʇ_, **Supplementary Figure S3**).

Recordings of a single TREK-2 channel in the high *P_O_* mode both in presence and absence of NFx are shown in **Figure 3**. Inspection of these recordings reveals two distinct effects of the drug at all membrane voltages: a dramatic reduction in *P_o_* along with a reduction in the single-channel current amplitude (γ).

**Figure 3.**
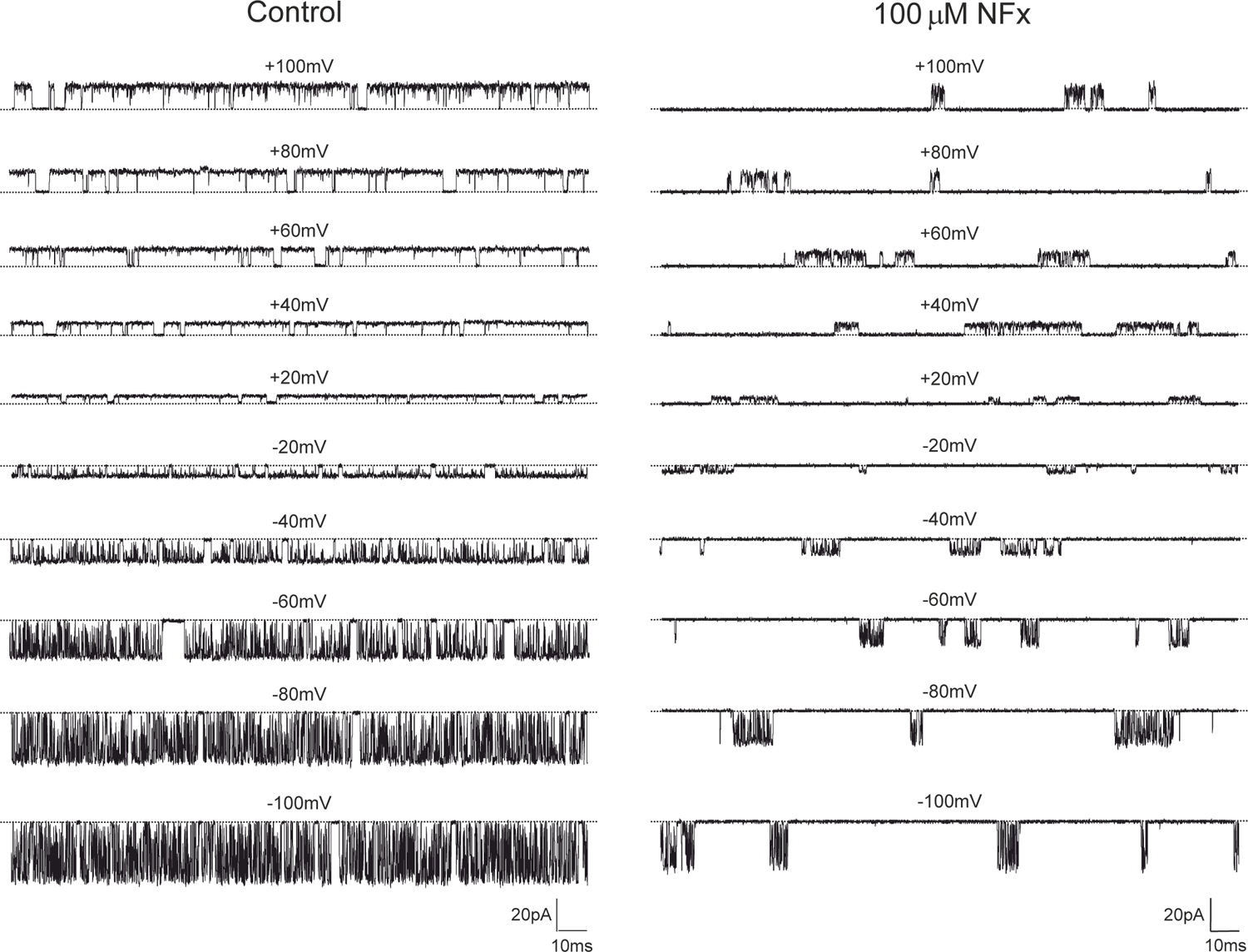
Effect of 100μM NFx on a single TREK-2 channel. Single-channel recordings of TREK-2 in in the high *P_o_* gating mode at different membrane voltages between +100mV and −100mV in the absence (left) and presence (right) of 100μM NFx. Dotted lines represent the closed channel levels.

The decrease in *P_o_* is associated with the appearance of very long closed periods that separate bursts of channel openings combined with brief closures. The reduction in γ induced by NFx was evident at both positive and negative voltages (**Figure 4A,B**), but as shown in **Figure 4B**, NFx also broadened the peak of the open current level reminiscent of classical open channel blocking mechanisms involving the fast binding and unbinding of blockers within a channel pore (Yellen, 1984). NFx could therefore also exert a combination of fast open channel block and an allosteric inhibition/inactivation effect on the selectivity gate similar to the effect of some ions on the filter gate of MthK channels (Thomson et al., 2014). However, other explanations for this reduction in γ are also possible and are examined later.

**Figure 4.**
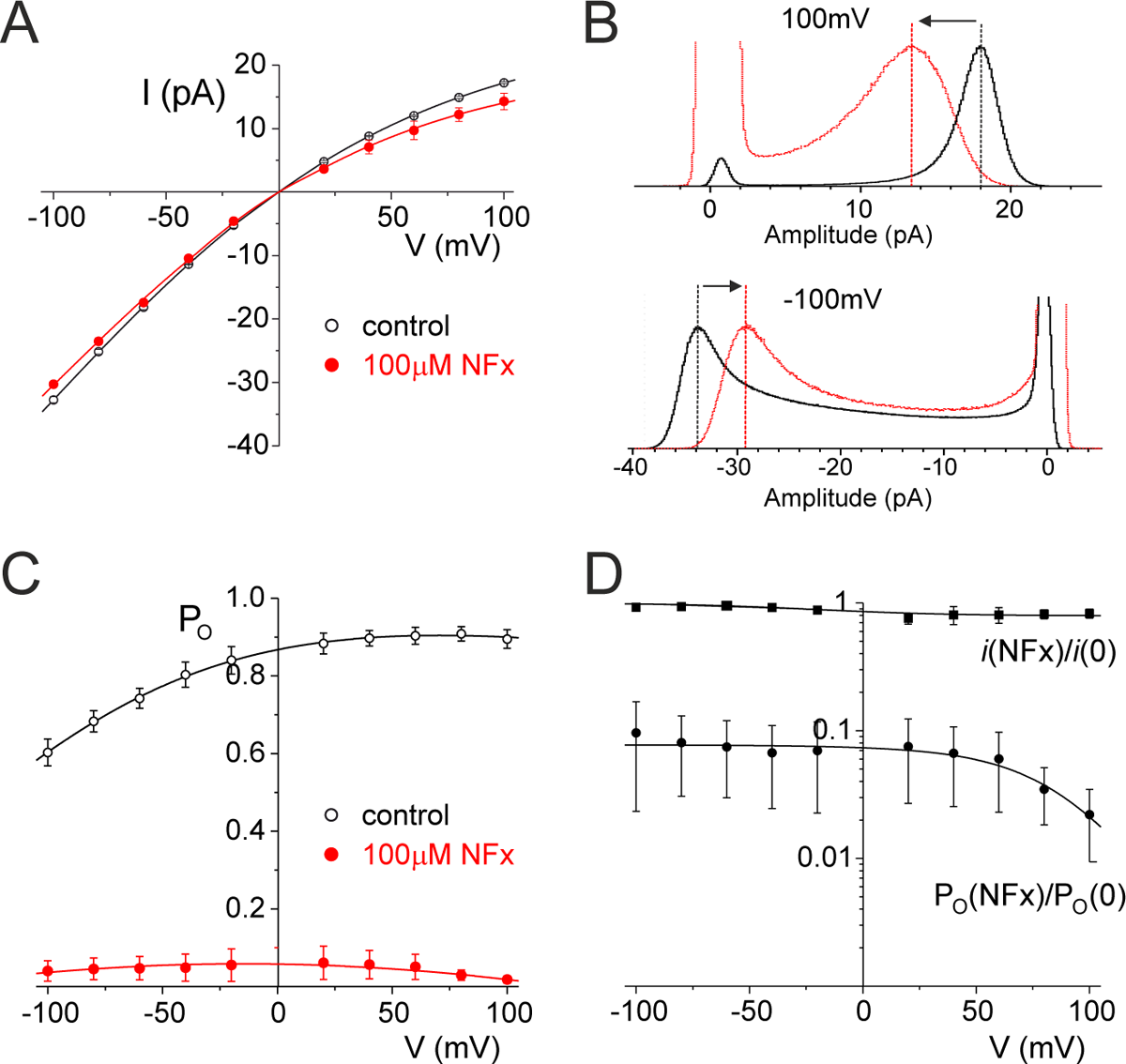
Summary of the effects of NFx on TREK-2 single channels. **A.** The mean single-channel conductance of TREK-2 in the absence (open circles; n=3) and presence of 100μM NFx (filled circles; n=3). **B.** Histograms of open single-channel current level in the absence (black line) and presence (grey line) of 100μM NFx at +100mV (top) and −100mV (bottom). **C.** Mean single-channel open probability of TREK-2 in the absence (open circles; n=3) and presence of 100μM NFx (filled circles; n=3). **D.** Mean single-channel current conductance (filled squares) and mean single-channel open probability (filled circles) in the presence of 100μM NFx normalized to values obtained in the absence of NFx (n=3).

### The effect of NFx on TREK-2 channel P_o_ is voltage-dependent

The inhibition of WT macroscopic TREK currents by NFx has previously been reported to be voltage-independent between +60 and −60 mV (Kennard et al., 2005; McClenaghan et al., 2016) and when we examined its effects on the *P_o_* of channels reconstituted in bilayers, we also found its inhibitory effects to be voltage-independent below +60 mV. However, when channel *P_o_* was measured above +60 mV, some voltage-dependence was observed, with increased efficacy at more depolarized potentials (**Figure 4D**).

To determine the relevance of these findings to full-length WT channels we re-examined the voltage-dependence of NFx inhibition of macroscopic WT TREK-2 currents expressed in *Xenopus* oocytes. Previous studies examined only a single NFx concentration which produces ∼80% inhibition (Kennard et al., 2005). We therefore determined macroscopic dose-response relationships for NFx inhibition at depolarized potentials and found a modest voltage-dependence with a slightly increased efficacy at more positive voltages (**Supplementary Figure S4**). Intriguingly, this finding may account for some of the minor variations in *IC_50_* values reported in the literature where inhibition was recorded at different potentials (Kennard et al., 2005; Dong et al., 2015; McClenaghan et al., 2016). It will therefore be important to take this voltage-dependent effect into consideration when reporting future *IC_50_* values for NFx inhibition.

### The effect on NFx is not state-dependent

The original model shown in **Figure 1A** implies two possible mechanisms of channel opening from either the down state or the up state. If the filter gate only opens when the channel is in the up conformation this might explain why many activators decrease NFx efficacy and why NFx can slow the kinetics of activation, but it does not explain why NFx inhibition is unchanged when the channel is activated by 2-APB or by Rb^+^. To determine if there was any state-dependence we therefore examined whether there was any correlation between channel *P_o_* and NFx efficacy, and whether there were any differential effects of NFx on the open and closed times of the channel.

We first determined whether NFx affects TREK-2 channels differently in the standard and high *P_O_* mode, but found no obvious difference even though their *P_o_* values differ markedly; at 10μM NFx the high *P_o_* mode was blocked by 20 ± 0.02% (n=3; mean *P_o_*=0.85±0.05) and the standard mode by 23 ± 0.05% (n=5; mean *P_o_*=0.40±0.05).

We next analyzed the effect of NFx on the kinetics of channel gating in the high *P_o_* mode. **Table 1** and **Figure 5A** shows that in the absence of NFx, the mean open time of the single apparent open state exhibits a strong bell-shaped dependence on membrane voltage with a maximum open time around +50 mV. It also shows a dramatic decrease in mean open time caused by NFx; although the effect was sharply reduced below −50 mV and virtually absent at −100 mV (**Figure 5E**).

**Figure 5.**
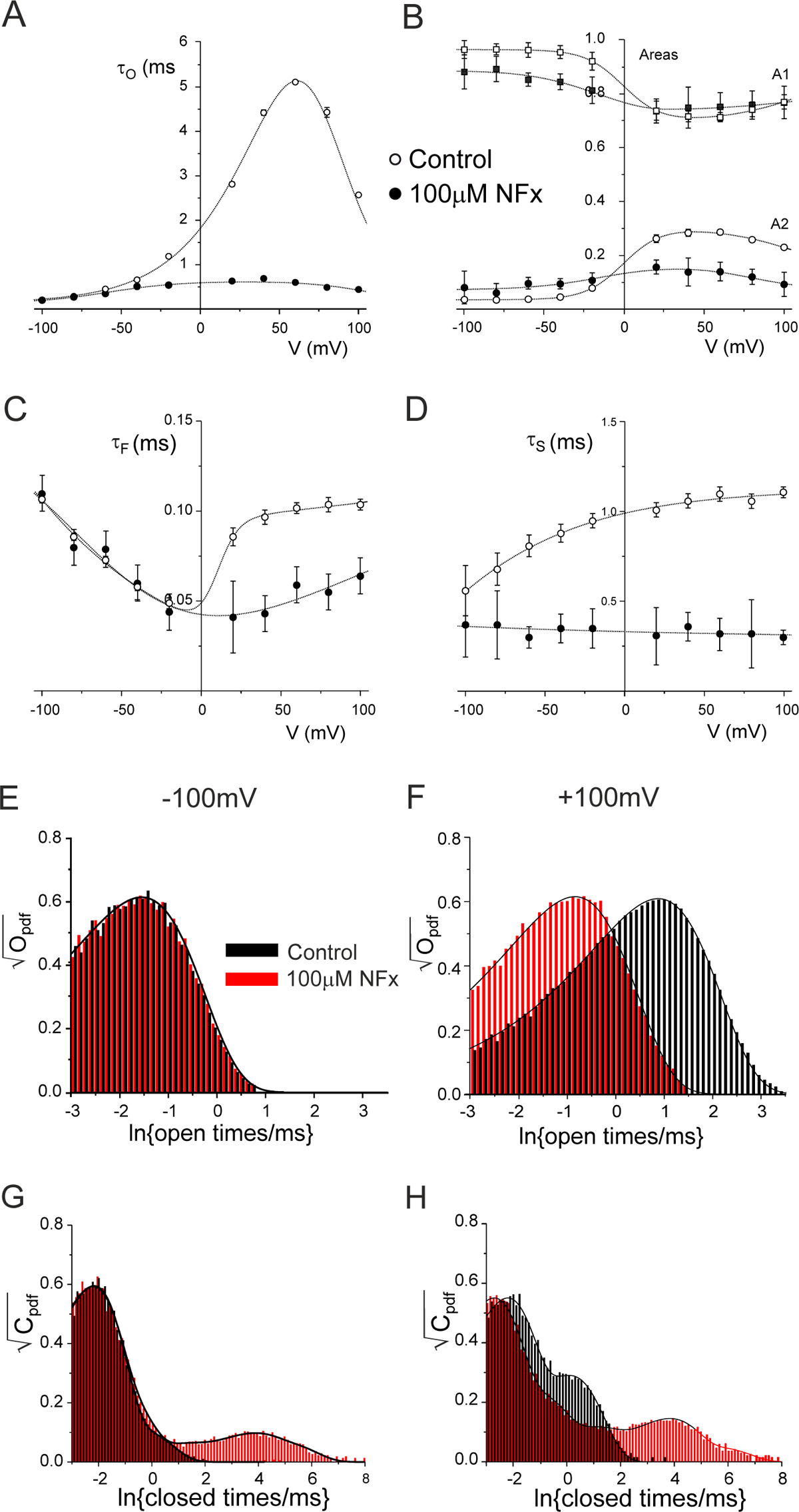
Effect of NFx on single-channel kinetics. **A.** Mean open time as a function of membrane voltage in the absence (open circles) and presence of 100μM NFx (filled circles) in single-channel recordings depicted in Figure 4. The lines through the data are drawn by hand. **B.** Relative areas of the shortest (squares) and the second shortest (circles) closed states in the absence (open symbols) and presence (filled symbols) of 100μM NFx. **C.** Mean shortest closed time as a function of membrane voltage in the absence (open circles) and presence of 100μM NFx (filled circles). **D.** Mean second shortest closed time as a function of membrane voltage in the absence (open circles) and presence of 100μM NFx (filled circles). **E.** Distribution of open times in the absence (open bars) and presence (filled bars) of 100μM NFx at −100mV. **F.** Distribution of open times in the absence (open bars) and presence (filled bars) of 100μM NFx at +100mV. **G.** Distribution of closed times in the absence (open bars) and presence (filled bars) of 100μM NFx at −100mV. **H.** Distribution of closed times in the absence (open bars) and presence (filled bars) of 100μM NFx at +100mV.

In the absence of NFx, the mean intrinsic short closed time (τ_F_) showed an inverted bell-shaped dependence on membrane voltage with a minimum around −30 mV (**Figure 5C, Table 1**). Similar to the effect of NFx on the mean open time, reduction of this closed lifetime by NFx was also suppressed at negative membrane voltages (**Figure 5C,G; Table 1**). In contrast to both mean open and short closed times, the mean long closed time in the absence of the drug (τ_S_) only exhibited a mild dependence on membrane voltage (**Figure 5D**). NFx reduced the mean lifetime of this component across the whole voltage range but its effects on the voltage dependence of the relative frequencies (areas) of the two intrinsic apparent closed states were more complex (**Figure 5B**).

Interestingly, NFx inhibition resulted in the appearance of three additional apparent closed state components (**Figure 5G,H; Table 1**). Their mean lifetimes showed no obvious voltage dependence, but the relative frequency of the two longest states (A_4_ and A_5_) appeared to increase with voltage **(Supplementary Figure S5A,B).** In particular, the steep increase in frequency of the longest closed state above +60 mV may account for the increased efficacy of NFx inhibition at more positive potentials. As shown in **Figure 3**, application of NFx on TREK-2 channel in the high *P_o_* mode resulted in a bursting behavior, with both mean burst and interburst durations exhibiting voltage-dependence above +60mV (**Supplementary Figure S5C,D**). Overall, these results suggest that NFx inhibition is not state-dependent and affects all intrinsic gating states of the channel.

To confirm that NFx inhibition of full-length WT TREK-2 channels is also state-independent (ie NFx affects not only closed but also open states of the channel), we examined its effects on the mean open time of TREK-2 channels expressed in HEK293 cells. As shown in **Supplementary Figure S6**, 10 μM NFx produced a clear reduction of τ_o_ of ∼30% thereby supporting the idea that NFx inhibition affects both the open and closed states.

The fact a large number of factors which change P_o_ are without effect on TREK-2 NFx sensitivity implies that the drug must bind with the same affinity to all open and closed states of the NFx-sensitive (down) conformation, and that NFx binding induces the same conformational changes in the filter gate to produce closure irrespective of whether the channel is closed or open. Single-channel analysis also revealed a dramatic increase in the number of apparent closed states in the presence of NFx (**Figure 5G,H**) suggesting the selectivity filter may alternate between several closed states *after* the conformational change(s) induced by NFx binding. However, the complexity of a kinetic scheme that would adequately describe such gating behavior combined with the relatively low accuracy with which the properties of very long closed states can be determined meant that we decided not to pursue this analysis of NFx-induced gating any further.

### What is the origin of the apparent voltage-dependence of NFx block?

The voltage-dependence of NFx inhibition is puzzling as there is no strong voltage-dependence to inhibition by other positively charged QA ions such as TPA and THexA which also bind deep within the inner pore, and there is no obvious intrinsic voltage-dependence of *P_o_* in the positive voltage range where this effect becomes apparent (**Figure 4C**).

To understand this, we modelled the voltage-dependent gating of TREK-2 in the high *P_o_* mode in the absence of NFx. Although the high *P_o_* mode could be well described by just three states (**Figure 5**), we found that 3-state kinetic models were unable to describe this voltage-dependence. Instead, it was necessary to assume that each of these three states were composites of several states whose distributions change dramatically with membrane voltage (**Supplementary Figure S7**). This kinetic model was capable of describing the voltage-dependence of single-channel parameters in the absence of NFx and is described in the **Supplementary Information**. The model predicted that the voltage-dependent filter gating above +60mV is dominated by a distinct set of kinetic states (O_3_, C_F3_ and C_S3_ in **Supplementary Figure S7C**). This behavior might therefore also be responsible for the voltage-dependent inhibitory effect of NFx at these positive voltages (Figure 4D).

Interestingly, the interaction of QA ions with the filter is also known to be capable of producing a range of voltage-dependent effects on filter gating in both CNG channels and MthK K^+^ channels (Martinez-Francois et al., 2009; Posson et al., 2013). It is therefore tempting to speculate that similar mechanisms may occur here.

### Electrostatic effects of NFx binding do not contribute to inhibition

Although our results suggest additional allosteric effects of NFx on the filter gate itself, the reduction in *γ* combined with the broadened peak of open current level (**Figure 4B**) are also reminiscent of a classical open channel blocking mechanism (Yellen, 1984). Furthermore, at physiological pH NFx is positively charged and its orientation within inner cavity would point these charged groups towards the permeation pathway (**Figure 1B**). We therefore decided to examine whether these positive charges directly affect K^+^ permeation and thereby contribute to the inhibitory effects of NFx on *P_o_* and/or *γ*.

We first used the crystal structure of TREK-2 obtained with NFx bound (PDB ID: 4XDK) and performed Poisson-Boltzmann electrostatic calculations for K^+^ along the axis of the pore in either the presence or absence of charged NFx. However, these calculations revealed only a modest increase in barrier to K^+^ permeation in the presence of NFx (**Figure 6A**).

**Figure 6.**
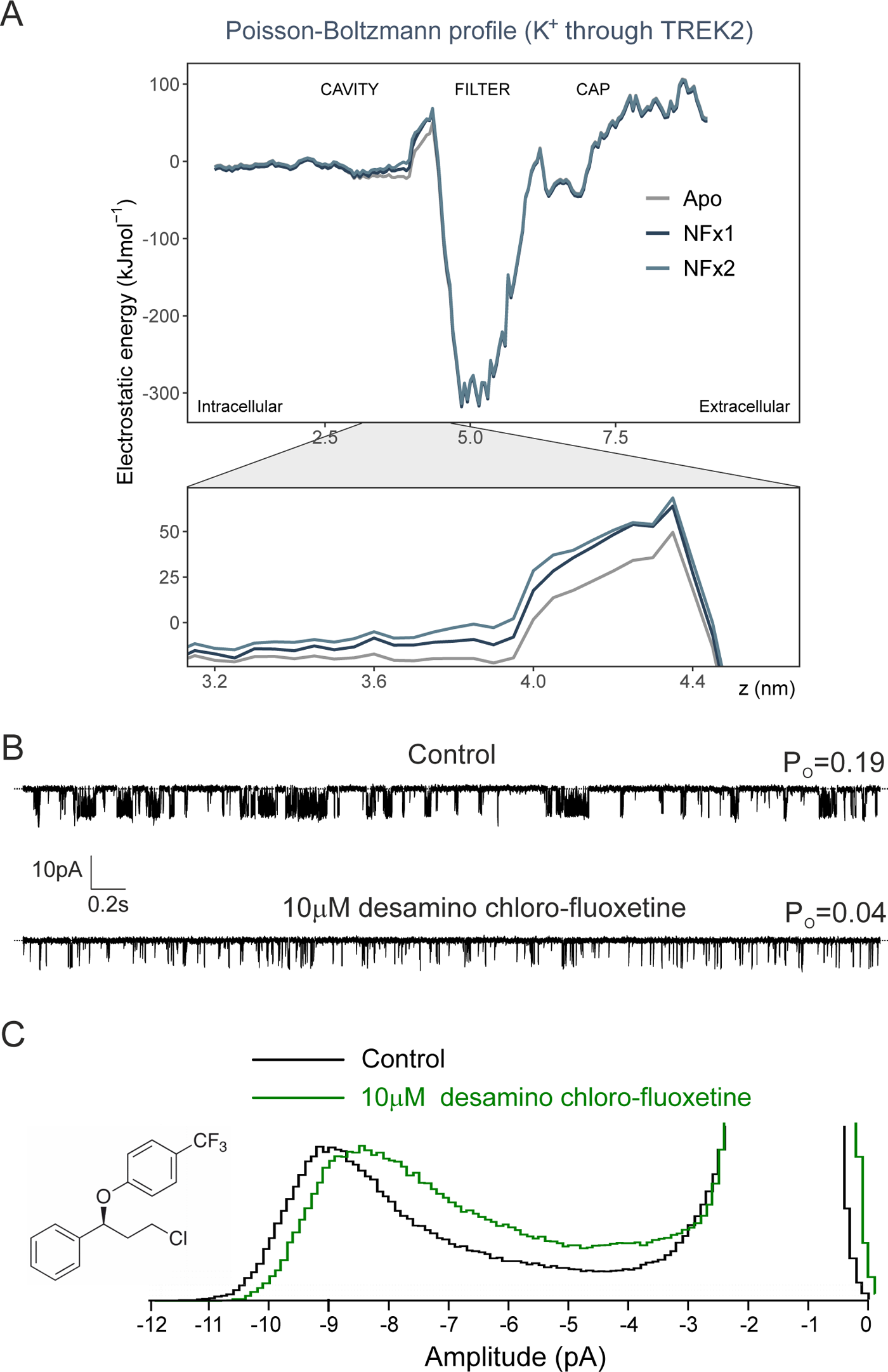
The relative charge of NFx does not contribute to its inhibitory effects. **A.** Electrostatic profile of a K^+^ through the pore of the TREK-2 channel in the presence and absence of NFx. The relevant region where NFx binds just below the filter is expanded below and shows two independent calculations in the presence of NFx compared to the Apo structure. A minor increase in the barrier for K^+^ permeation is seen in the presence of two charged NFx molecules bound at their sites below the filter. **B.** Single-channel recording of TREK-2 channel at −40mV in an excised patch from HEK293 cells either in the absence or presence of an uncharged NFx derivative (10μM desamino chloro-fluoxetine) as indicated. Dotted lines in top traces represent the closed channel level. **C.** Histograms of single channel currents from recordings shown above. For clarity, both amplitude histograms were scaled to the same open channel level. The chemical structure of desamino chloro-fluoxetine is shown on the left.

To examine the role of this charge on channel activity we next tested the inhibitory effect of a ‘neutralized’ NFx derivative on WT TREK-2 channels expressed in HEK293 cells. Desamino chloro-fluoxetine is a NFx derivative in which the positively charged −NH_3_ is substituted by chlorine, but as shown in **Figure 6B,C**, 10 μM desamino chloro-fluoxetine also markedly reduced channel *P_o_* (75 ± 0.05%; n=5) with a concomitant reduction in *γ*. (15 ± 5%; n=5). This effect was similar to that produced by charged NFx under identical conditions (70 ± 0.03% for *P_o_* and 11 ± 7% for γ, respectively n=5). Together these results demonstrate the charged nature of NFx contributes little to its inhibitory effects on channel activity or conductance.

### Impaired channel gating can produce dynamic changes in NFx sensitivity

The presence of NFx within its binding sites will clearly contribute to its inhibitory effects by stabilizing the channel in the down conformation, but our results also indicate allosteric effects on the filter gate itself. Interestingly, many channel regulators also operate via allosteric coupling of movement in the transmembrane helices to changes in the filter gate, and several activatory mutations have been shown to impair this coupling process (Brohawn et al., 2014; Lolicato et al., 2014; Dong et al., 2015; Zhuo et al., 2016). The reduced NFx efficacy seen in these activatory mutations may therefore result from altered allosteric coupling rather than changes in *P_o_*.

To explore this, we examined the effect of NFx on the single channel behavior of TREK-2 channel with the Y315A mutation located within the ‘hinge’ region on M4 which interacts with M3. This mutation markedly increases macroscopic TREK-2 currents and has been reported to reduce, but not abolish, NFx inhibition (McClenaghan et al., 2016).

WT and Y315A mutant channels were therefore expressed in HEK293 cells. As expected for WT TREK-2, only channels with a very low *Po* were observed (ranging 0.04 - 0.13 at −40 mV). However, unlike many other ‘NFx-insensitive’ activatory mutations which only produce a modest increase in *Po* (e.g. mutation of the ‘pH-sensor’ glutamate in the proximal C-terminus) (Bagriantsev et al., 2011), we observed a markedly higher *Po* (0.71 ± 0.1, n=3 at −40mV) for Y315A mutant channels, i.e. ∼10-20x greater than for WT TREK-2.

Interestingly, when we examined the effect of NFx on this mutation, we found that single Y315A currents underwent a rapid ‘desensitization’ to NFx inhibition. **Figure 7A** shows that although 10μM NFx initially reduced channel *Po* by more than 90%, within ∼30 seconds this inhibitory effect was dramatically reduced. For comparison, the *Po* of WT TREK-2 channel was reduced by 79 ± 0.05% (n=6) and did not change further in the presence of NFx, or during repetitive applications of the drug.

**Figure 7.**
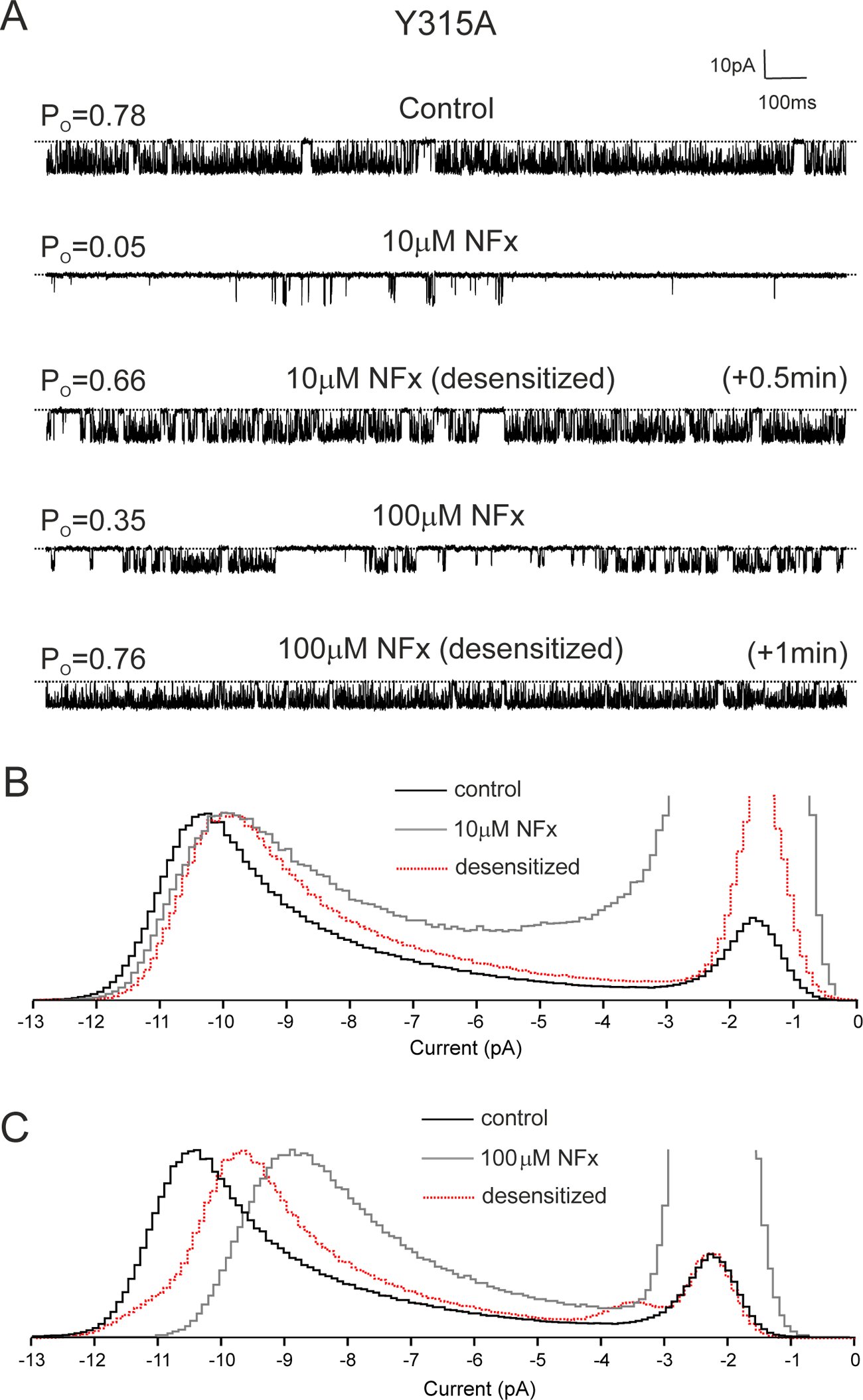
Desensitization of NFx Effects on TREK-2 Y315A. **A**. Single-channel recordings of TREK-2 Y315A mutant channels recorded at +40mV in excised patches from HEK293 cells in the presence and absence of either 10μM or 100μM NFx before and after channel desensitization. **B,C.** Histograms of single channel current amplitudes obtained in control solution and in the presence of 10μM or 100μM NFx before and after desensitization as indicated. For clarity, all amplitude histograms were scaled to the same open channel level.

Similar to its effect on WT TREK-2, NFx also reduced the single-channel current amplitude of Y315A TREK-2 and this effect was not abolished by desensitization (**Figure 7B**). Subsequent application of 100 μM NFx further decreased channel *PO*, but this effect also reversed within ∼1 minute so that channel *Po* before and after addition of 100 µM NFx were similar (**Figure 7A**). In addition to this relatively transient effect on channel *Po*, 100 μM NFx also reduced γ even further, but this was partially reversed by desensitization (**Figure 7C**).

The desensitization of this particular mutant to NFx could arise from several possible mechanisms; it could be caused by either abolished drug binding to the channel or a reduced ability of NFx to allosterically inhibit the channel via the filter gate itself. However, the fact NFx can still reduce γ even when its effect on *Po* is virtually abolished suggests the drug remains bound and that it is the allosteric effect of NFx on the filter gate which is impaired by this mutation.

Our results with uncharged NFx indicate that the reduction in γ is unlikely to represent pore block as the charge of the drug is not important. An alternative explanation for the reduction in γ could be that NFx stabilizes an ultrafast flickery closed state of the channel that results in a reduced apparent or measured γ rather than the true conductance itself.

### Allosteric antagonism of NFx inhibition by ML335

In **Figure 2** we show that activation by ML335 dramatically antagonizes the inhibitory effect of NFx on macroscopic TREK-2 currents. We therefore examined this antagonistic effect at the single-channel level on WT TREK-2 channels expressed in HEK293 cells. Interestingly, although 100 μM ML335 produces maximal activation at the macroscopic level it produced either partial or full activation of single TREK-2 channels (**Figure 8**). However, regardless of the resulting *Po* from either partial or maximal activation by ML335, subsequent application of 10 μM NFx failed to elicit any visible effects on either the mean *Po* (0.69±0.2, n=4 in both cases) or single channel current amplitude (the amplitude ratio of *γ* in both conditions was identical: 1.00 ± 0.01, n=4).

**Figure 8.**
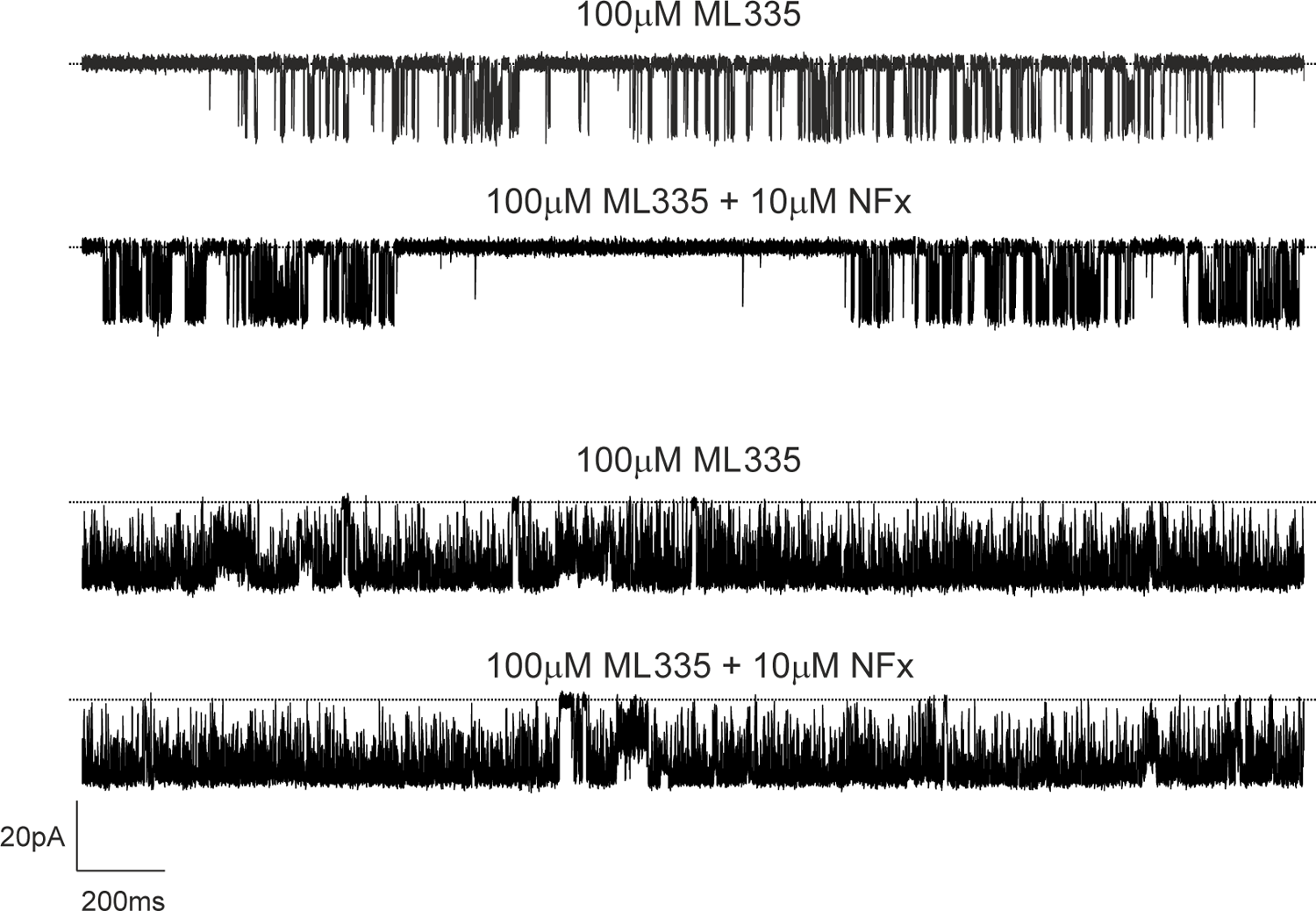
Norfluoxetine effects on single TREK-2 channel properties are abolished in the presence of 100μM ML335. Examples of single-channel recordings of TREK-2 channels at −40mV in excised patched from HEK 293 cells in the presence of 100μM ML335 having either partial (**top two traces**) or maximal (**bottom two traces**) effect on single-channel open probability either in the absence or presence of 10μM NFx, as indicated. Dotted lines in top traces represent the closed channel level. Note that NFx has no effect on either *P_o_* or γ in the presence of ML335.

The binding sites for these two drugs are distant from each other and given the apparent effect of ML335 on the conformational dynamics of the filter gate (Lolicato et al., 2020) the reduction in NFx efficacy is likely due to the fact that both mechanisms converge on the filter gate and prior activation of this gate by ML335 interferes with the transduction mechanism which couples NFx binding to this gating mechanism.

### Conclusions

The molecular mechanisms by which NFx inhibits TREK channels were previously unclear and impacted our ability to dissect the global structural movements underlying channel gating. By using a combination of macroscopic and single channel recordings we now provide clear evidence that NFx acts as a state-independent inhibitor which affects channel gating in several different ways.

Not only does NFx affect the equilibrium between the up and down conformations, but we now show that it also exerts state-independent allosteric control of the filter gate to influence both the open and closed states of the channel. We also show that 2-APB can robustly activate TREK-2 channels without affecting NFx sensitivity, thus demonstrating that the NFx-sensitive down conformation can also support a highly active open state. This also explains why not all activators impact NFx inhibition, something that would be impossible if opening of the filter gate only occurred from the up state where NFx cannot bind. These results allows us to expand the original gating scheme to include these different modes of gating and the effects of NFx on the filter gating mechanism (**Figure 9**).

**Figure 9.**
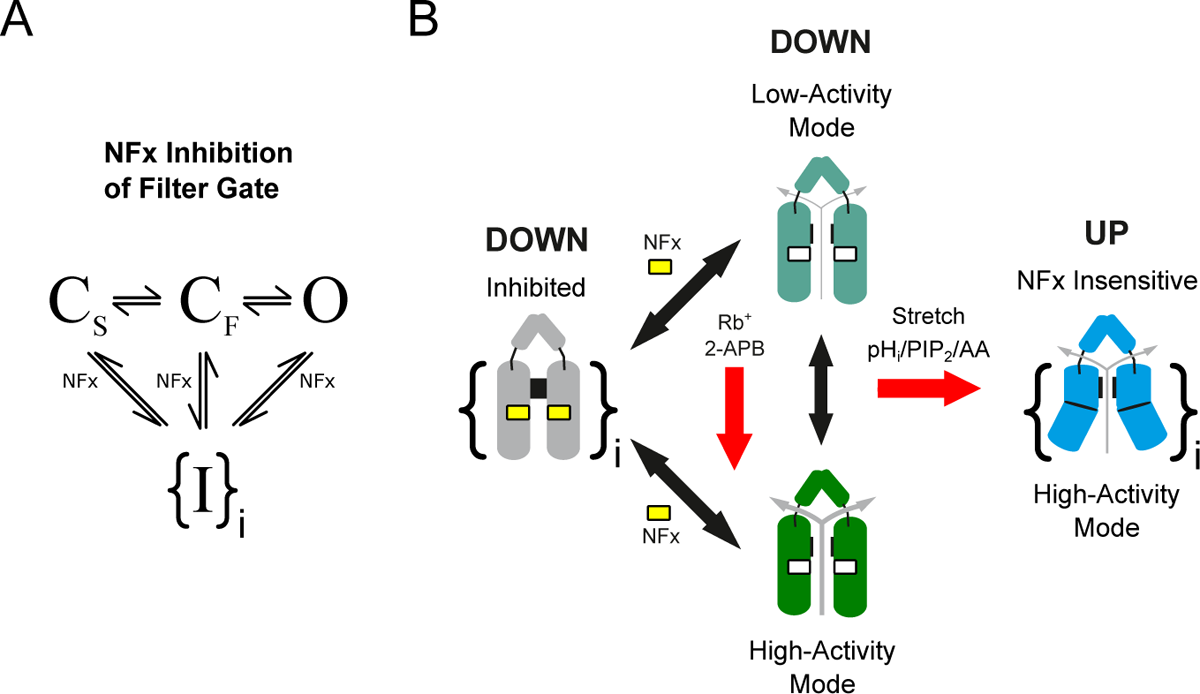
State-independent inhibition of TREK channels by NFx. **A.** Simplified filter gating scheme indicating that NFx interacts with both the open state as well as the long-lived (C_s_) and short-lived (C_F_) closed states of the filter gate to produce inhibition. The data suggest that multiple (i>1) inhibited states exist as indicated by the brackets. **B.** Summary cartoon indicating the different modes of channel behavior. NFx binding within the fenestrations prevents channels from moving into the up conformation, but we also now show that NFx inhibition affects both the open and closed states to produce multiple distinct closed states. TREK-2 can also adopt a high activity mode of gating in the down conformation e.g. when activated by 2-APB or Rb^+^ (red arrow). Other stimuli such as intracellular pH, PIP_2_ and membrane stretch (red arrow) are thought to promote high activity modes of gating by stabilization of various forms of the (NFx-insensitive) up conformation.

Our results also reveal a mild-voltage-dependence of NFx inhibition arising from an intrinsic voltage-dependent gating process within the selectivity filter which results in an increased efficacy of the drug at depolarized potentials. We also show that the reduction in single channel conductance produced by NFx results primarily from allosteric modulation of the filter gate rather than open pore block, and that the positive charge on the drug is not essential for its inhibitory effects. Overall, our results highlight the structurally divergent nature of the regulatory mechanisms which converge on the filter gate – a process which helps integrate inputs from a diverse range of physiological stimuli to effect polymodal regulation of TREK channel activity.

## Acknowledgements

We thank members of our labs for helpful comments on the manuscript and Mark Sansom for his support of the project. This work was supported by the BBSRC, the Wellcome Trust and by the Deutsche Forschungsgemeinschaft (DFG) as part of the Research Unit FOR2518, DynIon.

The authors declare no financial interests.

## Materials and Methods

### Preparation of TREK-2 containing Giant Unilamellar Vesicles

Human TREK-2 (*KCNK10*) ‘crystal construct’ protein (Gly67 to Glu340) was expressed and purified as previously described (Dong et al) with the exception that it was purified in 1% w/v n-Octyl-β-D-Glucoside, 0.1% w/v cholesteryl hemisuccinate. 1,2-diphytanoyl-sn-glycero-3-phosphocholine (DPhPC) was dissolved in chloroform to a concentration of 10 mM and stored at −20°C. The GUVs where then made by electroformation in a 1 M sorbitol solution using the Vesicle Prep Pro (Nanion Technologies, GmbH). Purified TREK-2 was then mixed with GUVs to a final concentration of ∼1 - 5 µg/ml and incubated overnight at 4 °C with 0.5mg/ml Bio-Beads (Bio-Rad) prior to use.

### Clones and Chemicals

Full length human TREK-2 isoform 3 (NP_612191) was used throughout this study and was subcloned into the pFAW vector for expression in oocytes. The truncated construct used to generate protein (TREK-2ΔN/ΔC) is identical to that used previously to obtain crystal structures (Dong et al., 2015); it contains a deletion of 71 residues at the N-terminus and 213 residues at the C-terminus. NFx and desamino chloro-fluoxetine (Toronto Research Chemicals). NFx was dissolved in DMSO and diluted to working concentrations on the day of experimenting (max final DMSO concentration was 0.3 %). Desamino chloro-fluoxetine was dissolved in chloroform and diluted to working concentrations on the day of experimenting (max final concentration was 0.01 %).

### Bilayer recordings and analysis

All electrophysiological recordings were performed with the Nanion Port-a-Patch system connected to an Axopatch 200B amplifier via a Digidata 1440A digitizer (Molecular Devices). Data was filtered at 5 kHz and recorded at a 200 kHz sampling rate with program Clampex (Molecular Devices). Experiments were carried out in symmetrical 200 mM KCl, 10 mM HEPES (pH 6.0 with KOH) solutions. Single-channel currents were idealized using 50% threshold criterion with program Clampfit (Molecular Devices) at an imposed resolution of 50μs. Only one open and one closed level was considered in the analysis – all subconductance states were neglected (these typically comprised less than 1% of open and closed level events of channels in the “high *P_O_*” mode. Analysis of amplitude and dwell-time distributions was carried out in Origin (OriginLab Corporation) and an in-house software written in Mathematica (Wolfram Technologies). Empirical correction for open times due to missed events was carried out as described previously by (Davies et al., 1992). Critical time for burst analysis was determined using Colquhoun and Sakmann criterion (Colquhoun and Sakmann, 1985).

### Electrostriction measurements

Membrane elasticity measurements were performed as described previously (Vitovic et al., 2013). Briefly, 1kHz sinewave with an amplitude of 100mV was applied to the membrane using a wave generator (Rigol DG821, Rigol technologies). Due to the non-linear dependence of membrane capacitance on the voltage V (C = C_0_ (1 + αV^2^), where C_0_ is the capacitance at V = 0 and α is the electrostriction coefficient), third current harmonic with frequency 3 kHz and amplitude A_3_ is generated in addition to the basic first current harmonic, A_1_ (frequency 1 kHz). The ratio of modulus of elasticity with and without NFx (E_ꞱNFX_/E_Ʇ_ (0)) is then given as A_3_(0)/A_3NFX_, where A_3_(0) and A_3NFX_ are amplitudes of third current harmonic frequencies in the absence and presence of NFx, respectively.

### Expression in Oocytes and HEK293 Cells

Oocytes were prepared for injection of mRNA by collagenase digestion followed by manual defolliculation and stored in ND96 solution which contained (in mM): 96 NaCl, 2 KCl, 1.8 CaCl_2_, 1 MgCl_2_, 10 HEPES, (pH 7.4) and was supplemented with 2.5 mM sodium pyruvate 50 µg/ml gentamycin, 50 µg/ml tetracycline, 50µg/ml ciprofloxacin and 100 µg/ml amikacin. Cells were injected with 1-4 ng of mRNA up to 4 days post isolation. In vitro transcription of mRNA was done using the Amplicap™ SP6 Kit, (Cambio). Experiments were performed 12-24 hours post injection at room temperature (22 °C unless otherwise indicated). For measurement and comparison of basal whole-cell currents oocytes were injected with 4 ng of RNA and recorded exactly 24 hours post injection.

HEK293 cells were cultured in DMEM (Sigma) containing 10% FBS (Life Technologies), 3 mM glucose, and 2 mM glutamine at 37°C in a humidified atmosphere of 5% CO_2_/95% O_2_ at 37°C. Cells were transiently transfected with 0.2 *μ*g of channel pcDNA3 per dish using FuGENE 6 according to the manufacturers’ instructions. Cells were used 1–2 days after transfection.

### Electrophysiology in heterologous systems and data analysis

For currents recorded in *Xenopus* oocytes: Giant-patch electrodes were pulled from thick-walled borosilicate glass and polished to give pipette resistances around 0.3-0.5 MΩ when filled with patch solution. Pipette solution contained (in mM) 116 NMDG, 4 KCl, 1 MgCl_2_, 3.6 CaCl_2_, 10 HEPES (pH 7.4); whilst bath solution contained (in mM) 120 KCl, 1 NaCl, 2 EGTA, 10 HEPES (pH 7.3). Patches were perfused via a gravity flow perfusion system. a HEKA EPC 10 USB single computer controlled amplifier and recorded using a Patchmaster v2×90.5 (HEKA electronics), filtered at 1 kHz and sampled at 10 kHz. For currents recorded in HEK293 cells: Patch electrodes were pulled from thick-walled borosilicate glass and polished to give pipette resistances around 3-5 MΩ when filled with patch solution. The currents were recorded from excised patches with both intracellular and extracellular solution containing 200 mM KCl, 10 mM HEPES (pH 6.0 with KOH). Patches were perfused via a gravity flow perfusion system. Data was acquired with pClamp and recorded using an Axopatch 200B (Molecular Devices), filtered at 5 kHz and sampled at 200 kHz. Single channel currents were analyzed in an identical manner to those obtained from bilayers (see above). The macroscopic concentration-inhibition relationships in Figures 1 & 2 were fitted with a modified Hill equation:

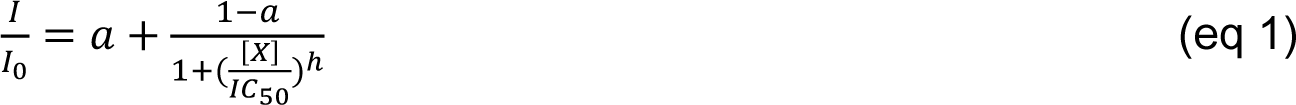

Where *I* and *I_O_* is the current in the presence and absence of inhibitor, *[X]* is the concentration of inhibitor, *IC_50_* is the inhibitor concentration at which the inhibition is half maximal, *h* is the Hill coefficient and *a* is the fraction of unblockable current; *a*=0 except where indicated in legend to Figure 1.

The macroscopic concentration-activation relationships in Figure 1E were fitted with a modified Hill equation:

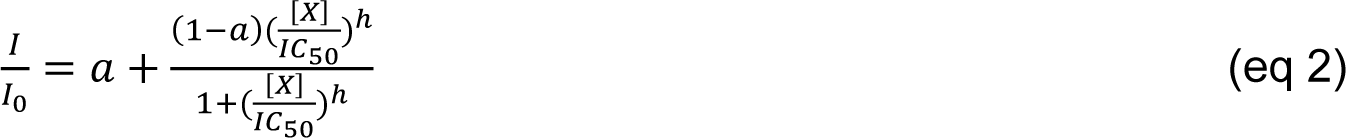

Where *I* and *I_O_* is the current in the presence and absence of agonist, *[X]* is the concentration of agonist, *IC_50_* is the agonist concentration at which the activation is half maximal, *h* is the Hill coefficient and *a* is the fraction of initial current.

### Poisson-Boltzmann electrostatics

Born energy profiles of a K^+^ ion through the channel pore of a TREK-2 structure (PDB ID: 4XDK) in the absence or presence of NFx, were calculated by numerically solving the linearized Poisson-Boltzmann equation using the Adaptive Poisson-Boltzmann Solver (Baker et al., 2001; Jurrus et al., 2018). The ion was positioned along the central channel axis at 0.05 nm intervals, extending 2 nm into the bulk phase from either side of the channel. A Born radius of 0.22 nm was used for K^+^. Protein and ligand atoms were assigned radii and partial charges from the CHARMM36 force field. The radius of an implicit solvent molecule was set to 0.14 nm, the ionic strength to 0.15 M KCl, and the dielectric constant to 78.5 for the solvent and 2 for the protein. The Born energy for inserting an ion at each sampled position was calculated at 37 °C by subtracting the individual electrostatic energies of the protein and the ion in solution from the electrostatic energy of the protein-ion system (Beckstein et al., 2004).

## SUPPLEMENTARY INFORMATION

### Notes on modelling

Each rate constant in the kinetic scheme in Figure S6C is a function of voltage, i.e:

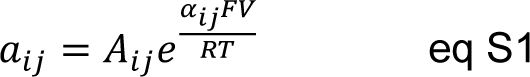

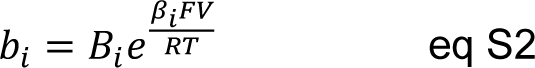

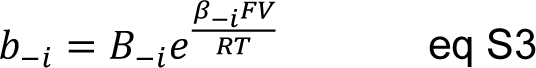

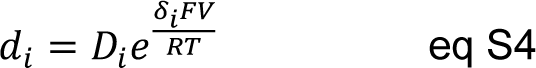

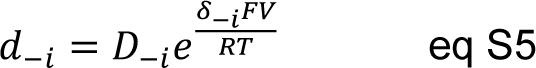

where indexes 1≤*i,j*≤ 3, *V* is the membrane voltage, *F* is the Faraday constant, *R* is the gas constant and *T* is the temperature. The Scheme in Fig 6C can be fitted either to dwell time distributions or, since it can be solved analytically, to the parameters of open and closed time distributions (state lifetimes and their corresponding areas).

Below −40mV, the data was best fit with a linear scheme O_1_-C_F1_-C_S1_ which forms the left part of the scheme in Supplementary Figure 6C with the following parameters:

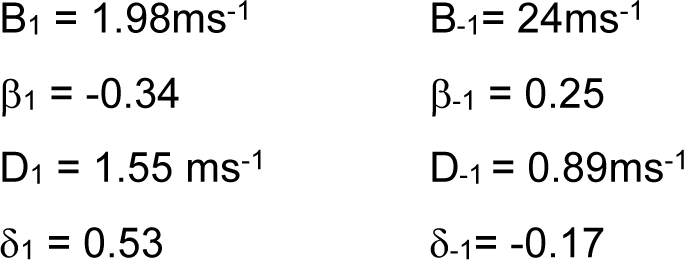

It was possible to reproduce the voltage dependence of data between −40mV and +60mV by adding states O_2_-C_F2_-C_S2_ with a connection between the two open states (connecting new states via closed states produced greater errors in fits). Finally, another set of three states O_3_-C_F3_-C_S3_ was necessary to add in order account for changes in voltage-dependence of open times and areas of closed times above +60mV. Models created by connecting various combinations of O-C_F_-C_S_ and C_F_-O-C_S_ states produced either poor fits of data of failed to converge to a solution. Analysis of the data with the kinetic scheme in Figure S6C revealed that there are many possible solutions which fit the data with similar accuracy with wide ranging values, particularly for constants A_ij_ and α_ij_. One of the solutions is shown in **Supplementary Figure S7D-G**.

The third set of kinetic states of the model (O_3_, C_F3_ and C_S3_) dominates gating above +60mV – a region where voltage dependence of NFx block has been observed (**Figure 4D**). This suggests that the drug’s voltage dependent effect can arise from the voltage-dependent intrinsic gating at the selectivity filter.

---

**Supplementary Figure S1:**
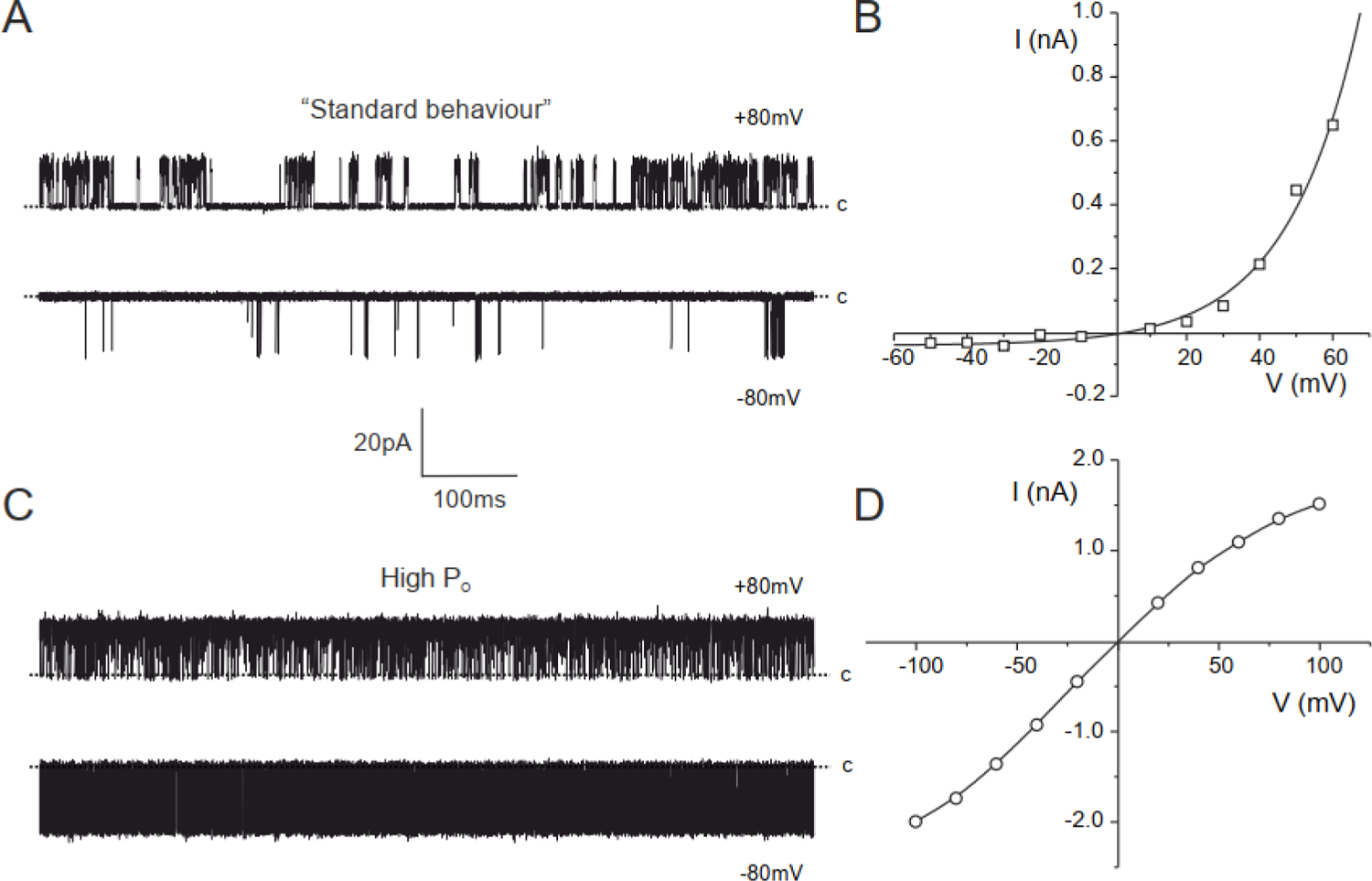
Two types of TREK-2 behavior in lipid bilayers. **A,B.** Single-channel recordings of TREK-2 incorporated into a bilayer at +80mV (top trace) and −80mV (bottom trace). The dotted line represents the closed channel level. **C,D.** Macroscopic current −voltage relationships simulated for one hundred TREK-2 channels using values of single-channel open probability (*P_O_*) and single-channel current amplitude (*i*) obtained from single-channel recordings of TREK-2 with standard (**C**) and “high *P_O_*” behavior (**D**). The lines are fit by hand.

**Supplementary Figure S2:**
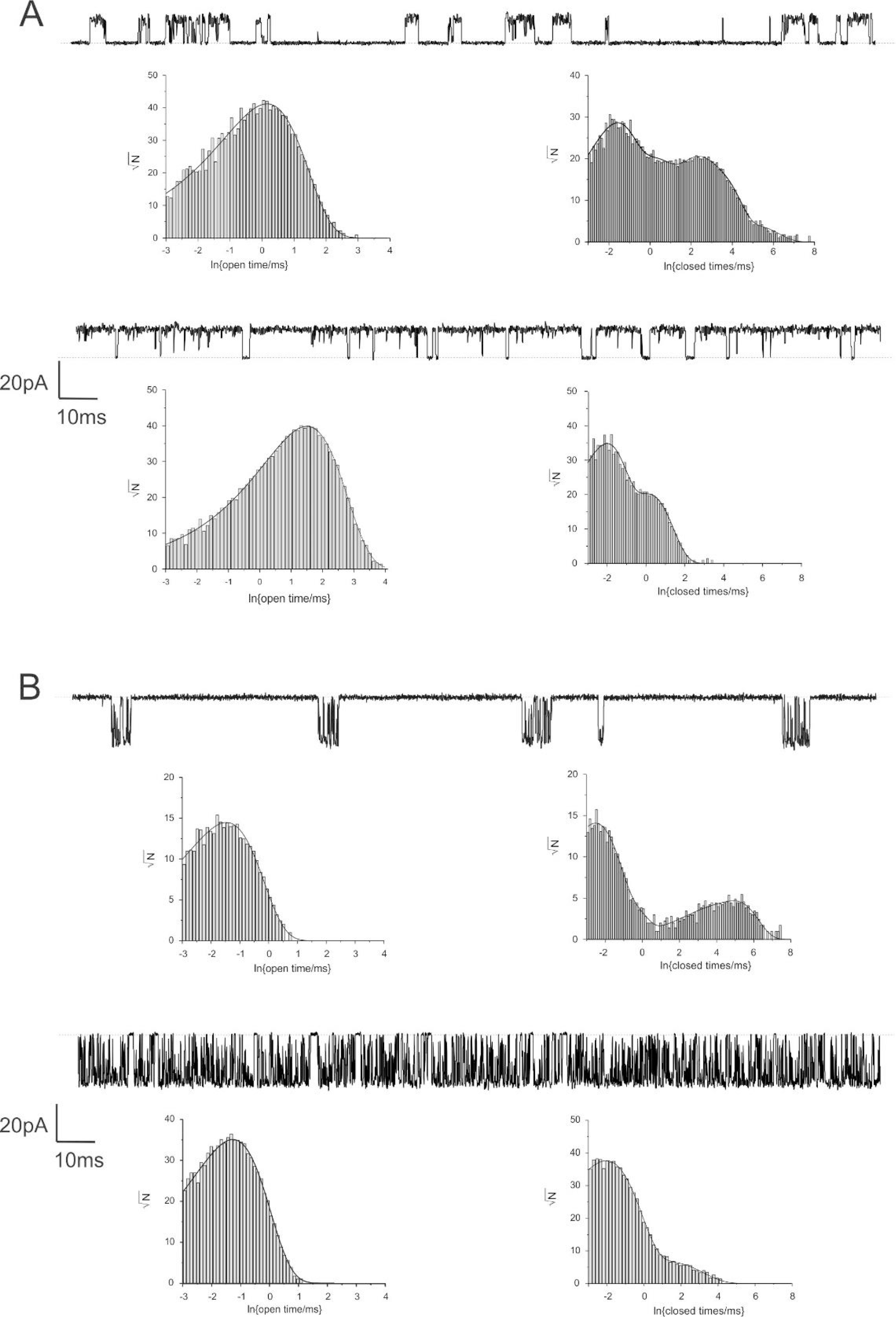
Comparison of single channel kinetics of TREK channels in standard and high *P_O_* mode. A. Outward currents. Top traces: single-channel recordings of TREK-2 in the standard (top trace, *P_O_* =0.14) and high *P_O_* mode (bottom trace, *P_O_* =0.92) at +60mV. Dotted line represents the closed channel level. Bottom panels: distributions of single-channel openings (left) and closures (right) obtained from recordings of TREK-2 in standard and “high *P_O_*” mode at +60mV. **Inward currents.** Top traces: single-channel recordings of TREK-2 in standard (top trace, *P_O_* =0.013) and “high *P_O_*” mode (bottom trace, *P_O_* =0.70) at −60mV. Dotted line represents the closed channel level. Bottom panels: distributions of single-channel openings (left) and closures (right) obtained from recordings of TREK-2 in standard and “high *P_O_*” mode at −60mV.

**Supplementary Figure S3.**
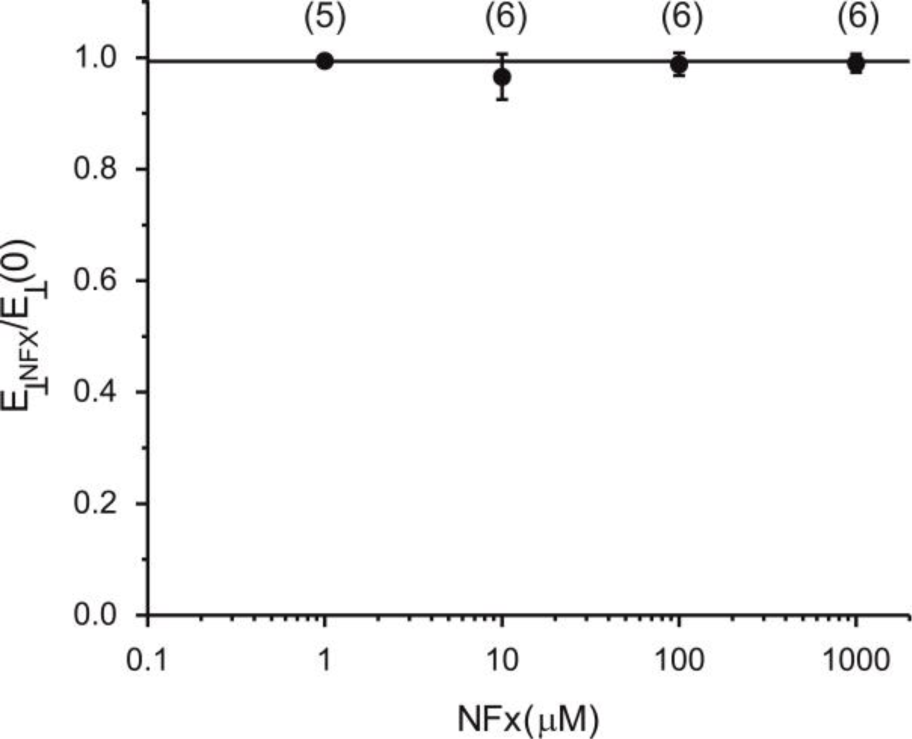
Effects of NFx on the physical properties of the membrane. The relationship between the modulus of elasticity in the perpendicular direction of the membrane in the presence of norfluoxetine (*E_ꞱNFX_*), normalized to that in control solution (*E_Ʇ_(0)*). Number of experimental values are shown above each point. The line is fit by hand.

**Supplementary Figure S4.**
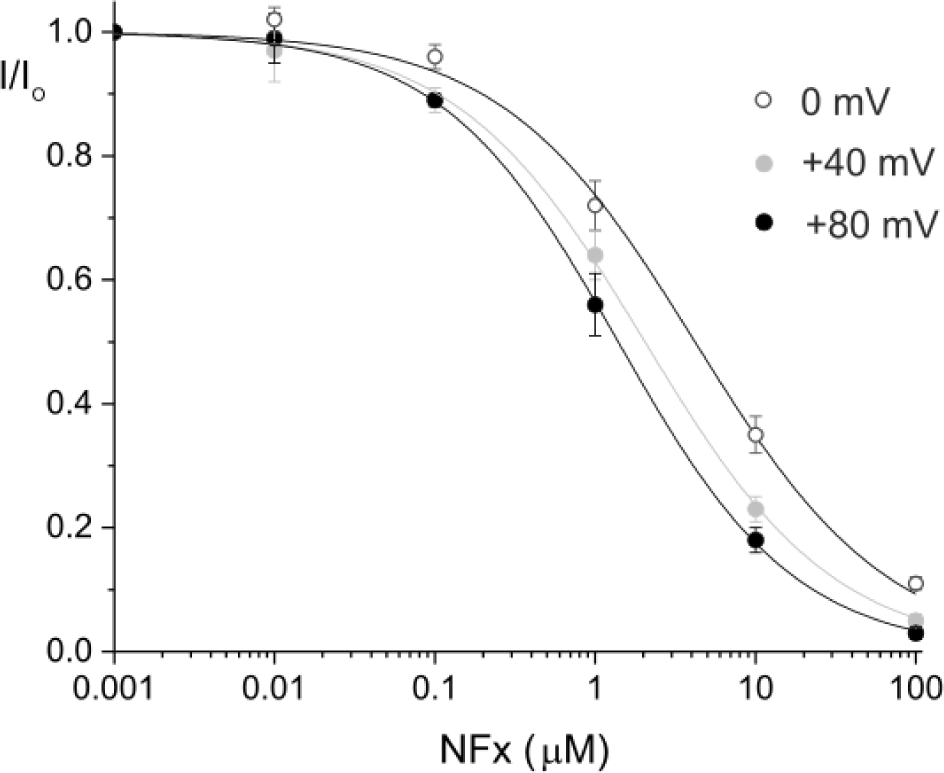
Voltage-dependence of NFx block at depolarized potentials. Doseresponse relationships were determined at different voltages for NFx inhibition of macroscopic currents in giant excised patches from oocytes expressing WT TREK-2. At saturating concentrations, relatively little voltage-dependence is observed but at depolarized potentials a small shift is observed. The lines are fit with a Hill inhibition equation (eq 1) assuming *a*=0. The *IC_50_* values are: 4.2 μM, *h*=0.72 (0 mV); 2.0 μM *h*=0.74 (+40mV); and 1.4 μM *h*=0.79 (+80mV).

**Supplementary Figure S5.**
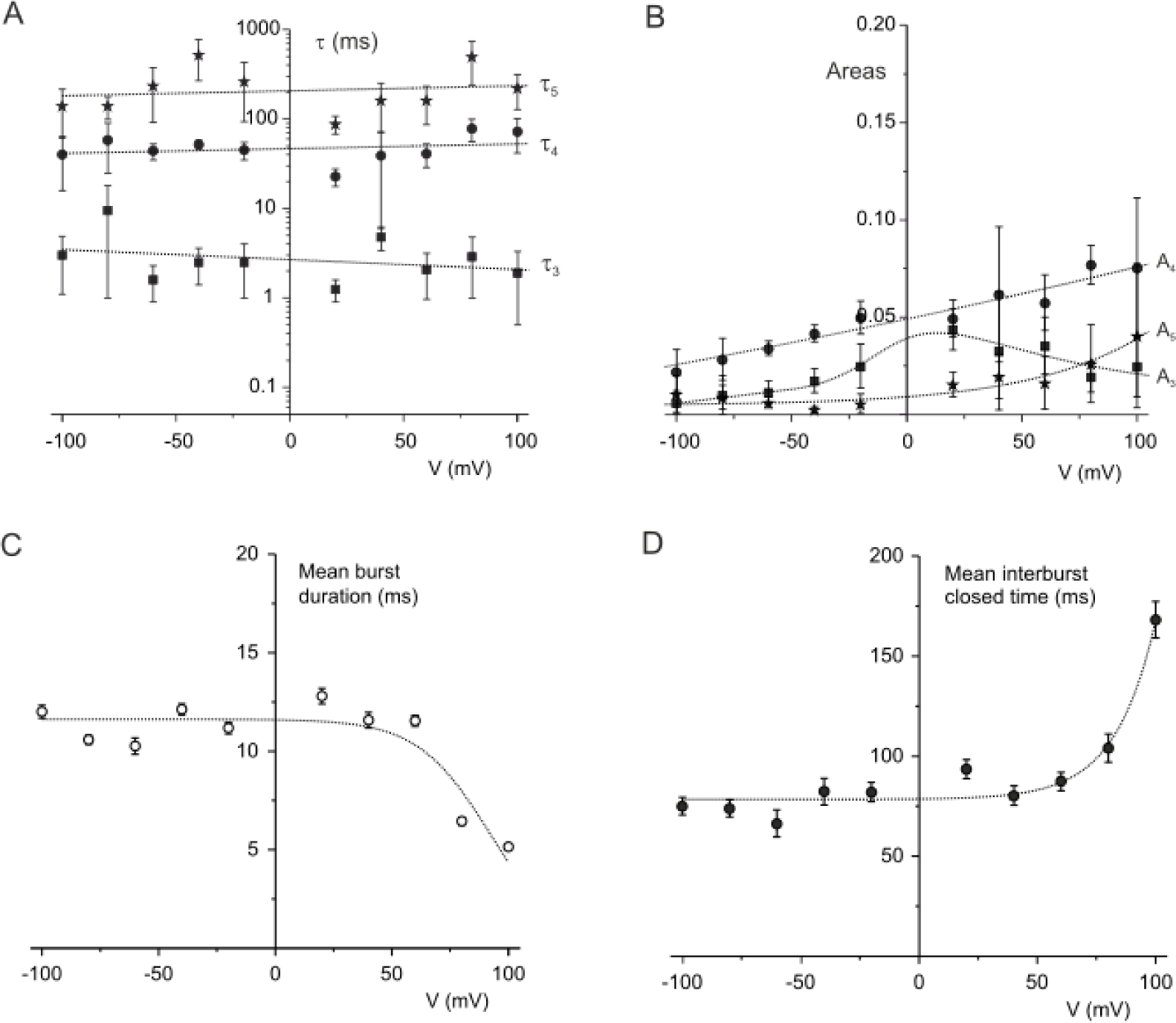
Properties of long closed states and bursts in the presence of NFx. Mean lifetimes (**A**) and relative areas (**B**) of three apparent long closed states observed in the presence of 100μM NFx in single channel recordings depicted in Figure 3. The lines through the data are fit by hand. **C,D.** The dependence of the mean burst duration and the mean inter-burst closure on the membrane voltage in the presence of 100μM NFx. This shows that the increase in NFx inhibition above +60 mV is accompanied by both a decrease in the mean burst length and an increase in the mean interburst close time. The lines are fit by hand.

**Supplementary Figure S6.**
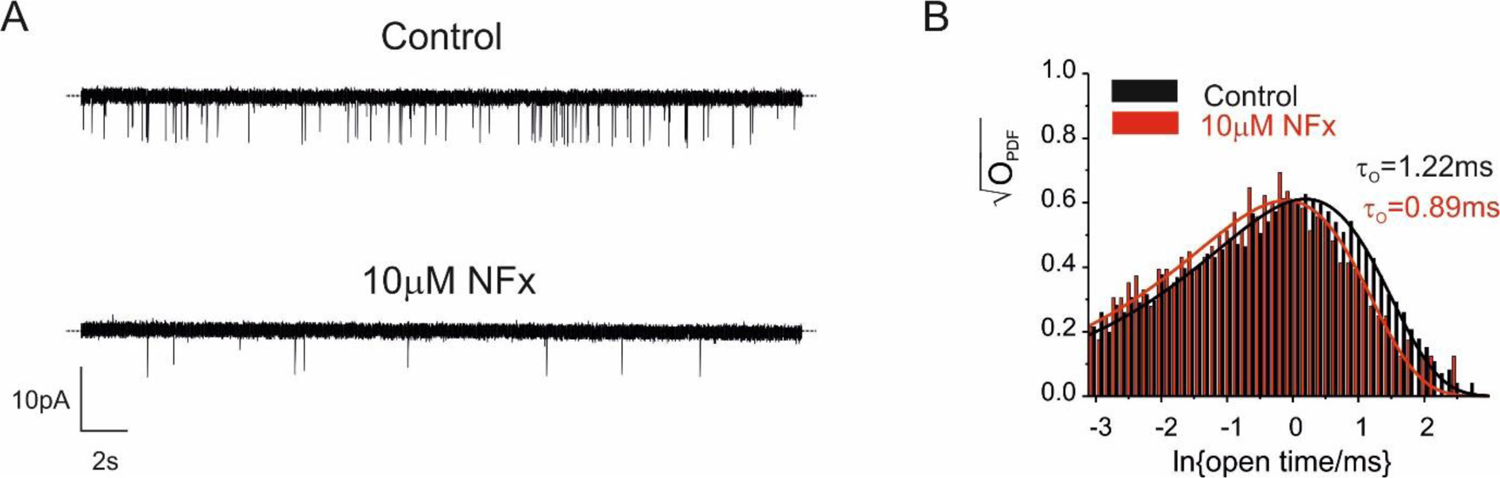
NFx affects open state of WT TREK2 channels expressed in HEK cells. **A**: Single-channel recordings of TREK-2 in the excised patch at −10mV in the absence (top trace) and presence of 10μM NFx (bottom trace). The dotted line represents zero current level. **B.** Dwell-time distributions of channel openings in the absence (black bars) and presence (red bars) of 10μM NFx. The lines are the best fit of the data to a single exponential function.

**Supplementary Figure S7.**
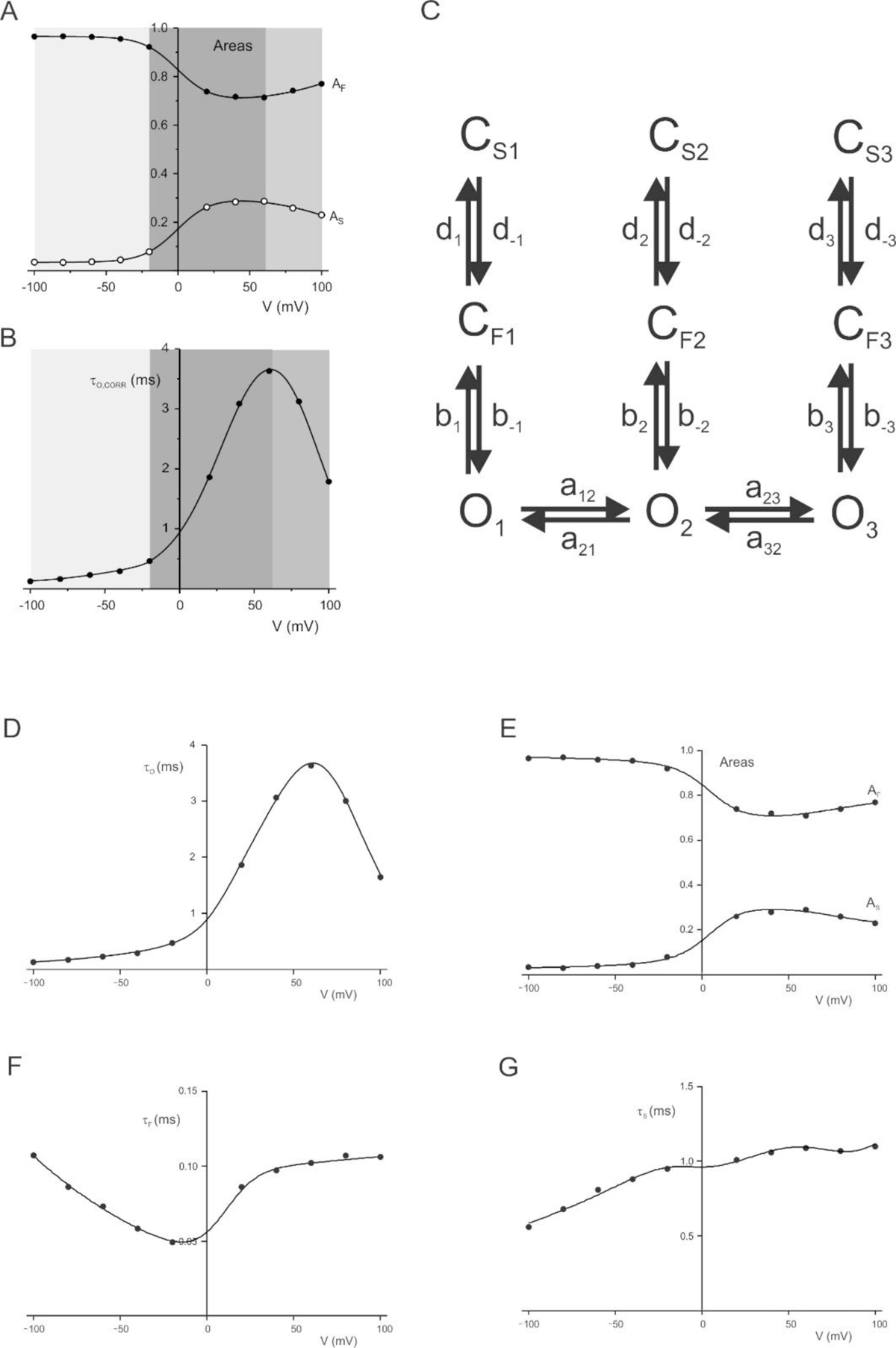
Kinetic model of gating at the selectivity filter. A. Voltage-dependence of relative areas of fast and long closed states. Three distinct regions characterized by different voltage-dependence are depicted in different shades of grey. **B.** Voltage-dependence of mean open times, corrected for missed events. Three distinct regions of behavior at these different voltages are highlighted in shades of grey. **C.** A kinetic scheme of the TREK-2 selectivity filter gate with three sets of open (O), short (C_F_) and long closed states (C_S_) affecting three distinct voltage regions depicted in panels A and B. **E-H.** Voltage-dependence of intrinsic mean open time **(E)**, short closed time **(F)**, long closed time **(G)** and relative areas of short and long closed times **(H)**. The lines are a fit of the kinetic model in Panel **C** to the data shown in Figure 5 with the following parameters: A_12_=5ms^-1^, A_21_=20 ms^-1^, α_12_=0.46, α_21_=-0.46, A_23_=21 ms^-1^, α_23_=0.95, A_32_=3.5 ms^-1^, α_32_=-0.95, B_2_=4.2ms^-1^, β_2_=1.3, B_-2_=7.2 ms^-1^, β_-2_=1.6*10^-5^, B_3_=6.2*10^-3^ms^-1^, β_3_=1.1, B_-3_=7.5, β_-3_= 2.1*10^-5^, D_2_=2.4, δ_2_ =1.2*10^-5^, D_-2_=1.6, δ_-2_=-0.17, D_3_=1.7ms^-1^, δ_3_=6.4*10^-5^, D_-3_=4ms^-1^, δ_-3_=-0.30.

**Table S1.** **Comparison of single-channel parameters of single TREK-2 channels depicted in Supplementary Figure S2**. τ_O_, mean open time; τ_Ci_ and A_Ci_, mean lifetimes and corresponding areas of closed times (*i*=1-5).

| Mode | V<br>(mV) | $\tau_o$<br>(ms) | $\tau_{C1}$<br>(ms) | $A_{C1}$<br>(%) | $\tau_{C2}$<br>(ms) | $A_{C2}$<br>(%) | $\tau_{C3}$<br>(ms) | $A_{C3}$<br>(%) | $\tau_{C4}$<br>(ms) | $A_{C4}$<br>(%) | $\tau_{C5}$<br>(ms) | $A_{C5}$<br>(%) |
| --- | --- | --- | --- | --- | --- | --- | --- | --- | --- | --- | --- | --- |
| Standard | +60 | 1.2 | 0.17 | 45 | 0.88 | 20 | 6.9 | 16 | 24 | 18 | 145 | 1 |
| High $P_o$ | +60 | 4.5 | 0.12 | 73 | 1.1 | 27 | - | - | - | - | - | - |
| Standard | -60 | 0.24 | 0.042 | 24 | 0.11 | 58 | 0.4 | 7 | 23 | 2 | 140 | 8 |
| High $P_o$ | -60 | 0.27 | 0.09 | 97 | 0.7 | 3 | - | - | - | - | - | - |

